# High-throughput discovery and characterization of human transcriptional effectors

**DOI:** 10.1101/2020.09.09.288324

**Authors:** Josh Tycko, Nicole DelRosso, Gaelen T. Hess, Aradhana, Abhimanyu Banerjee, Aditya Mukund, Mike V. Van, Braeden K. Ego, David Yao, Kaitlyn Spees, Peter Suzuki, Georgi K. Marinov, Anshul Kundaje, Michael C. Bassik, Lacramioara Bintu

## Abstract

Thousands of proteins localize to the nucleus; however, it remains unclear which contain transcriptional effectors. Here, we develop HT-recruit - a pooled assay where protein libraries are recruited to a reporter, and their transcriptional effects are measured by sequencing. Using this approach, we measure gene silencing and activation for thousands of domains. We find a relationship between repressor function and evolutionary age for the KRAB domains, discover Homeodomain repressor strength is collinear with *Hox* genetic organization, and identify activities for several Domains of Unknown Function. Deep mutational scanning of the CRISPRi KRAB maps the co-repressor binding surface and identifies substitutions that improve stability/silencing. By tiling 238 proteins, we find repressors as short as 10 amino acids. Finally, we report new activator domains, including a divergent KRAB. Together, these results provide a resource of 600 human proteins containing effectors and demonstrate a scalable strategy for assigning functions to protein domains.

## Introduction

In order to understand the molecular underpinnings of human biology and disease, we require systematic knowledge of how proteins function in human cells. Functional studies are practically challenging to approach systematically because the sequence space of proteins and their mutant variants is vast. For example, the human proteome likely includes over 1,600 transcription factors (Lambert et al., 2018). Extensive mapping efforts have generated atlases of the genome occupancy of a subset of these TFs. (ENCODE Project Consortium et al., 2020; Partridge et al., 2020). However, we still lack a complete understanding of their transcriptional effector functions. With systematic knowledge of which proteins can activate or repress transcription, which protein domains mediate these functions, and which residues in these domains are critical, we could build better models of gene regulation and devise better approaches to correct dysregulated gene expression in disease contexts. Thus, a scalable approach to determine and measure transcriptional effector functions is needed.

Traditionally, effector function is measured with a recruitment assay, in which a candidate effector protein is fused onto a synthetic DNA binding domain and recruited to a reporter gene promoter, resulting in a measurable impact on reporter gene expression (Sadowski et al., 1988). This approach is quantitative, and we have recently adapted it to measure the dynamics of transcriptional regulation in single mammalian cells using time-lapse imaging, revealing that transcriptional repressors from diverse KRAB, Polycomb, and HDAC complexes silence a reporter gene with distinct dynamics (Bintu et al., 2016). The recruitment assay has been applied extensively to characterize individual proteins and some fusions of multiple effectors (Amabile et al., 2016; Konermann et al., 2014), but the throughput of this assay is limited because each effector protein is individually cloned, delivered into cells, and measured. Recently, systematic recruitment of hundreds of chromatin regulators was achieved in yeast in an arrayed screen (Keung et al., 2014), and, in the last few years, pooled strategies for recruitment assays of activator domains were fruitfully implemented in yeast (Erijman et al., 2020; Staller et al., 2018) and Drosophila cells (Arnold et al., 2018).

Here, we report the development of HT-recruit, a high-throughput recruitment assay in human cells that allows us to measure the function of tens of thousands of candidate effector domains in parallel. The method combines pooled synthesis of oligonucleotide libraries encoding protein domains and variants, a synthetic surface marker reporter allowing facile magnetic cell separation, and next generation sequencing of the protein domains as a readout. This approach enabled us to quantify the effector activity of thousands of Pfam-annotated domains in nuclear-localized human proteins, providing a comprehensive functional assessment of large domain families of known relevance such as the KRAB and Homeodomain families. We also assign effector function to a number of Domains of Unknown Function (DUF). Further, we performed a deep mutational scan of the KRAB repressor domain used in CRISPRi (Gilbert et al., 2014), identifying mutations that ablate and enhance its gene silencing effects. Lastly, we identify compact repressor domains in unannotated regions of nuclear proteins using a library tiling 238 nuclear proteins. Together, we demonstrate a method for systematic functional analysis of effector domains, establish a resource for interpreting the roles of proteins in the human nucleus based on the capacity of their domains to activate or repress transcription, and generate a toolbox of compact and efficient effector domains to enable synthetic biology approaches to perturb and manipulate the epigenome and transcription.

## Results

### HT-recruit identifies hundreds of repressor domains in human proteins

In order to turn the classical recruitment reporter assay into a high-throughput assay of transcriptional domains, we had to solve two problems: (1) modify the reporter to make it compatible with rapid screening of libraries of tens of thousands of domains, and (2) devise a strategy to generate a library of candidate effector domains. To improve on our previously published fluorescent reporter (Bintu et al., 2016), we engineered a synthetic surface marker to enable facile magnetic separation of large numbers of cells and integrated the reporter in a suspension cell line that is amenable to cell culture in large volume spinner flasks. Specifically, we generated K562 reporter cells with 9xTetO binding sites upstream of a strong constitutive pEF1a promoter that drives expression of a two-part reporter consisting of a synthetic surface marker (the human IgG1 Fc region linked to an Igκ leader and PDGFRβ transmembrane domain) and a fluorescent citrine protein (**Figure 1A**). We used flow cytometry to confirm that recruitment of a known repressor domain, the KRAB domain from the zinc finger transcription factor ZNF10, at the TetO sites silences this reporter in a doxycycline-dependent manner within 5 days (**Figures S1A and S1B**). We also confirmed that magnetic separation with ProG Dynabeads that bind to the synthetic surface marker can separate cells with the reporter ON from OFF cells (**Figure S1C**).

**Figure 1.**
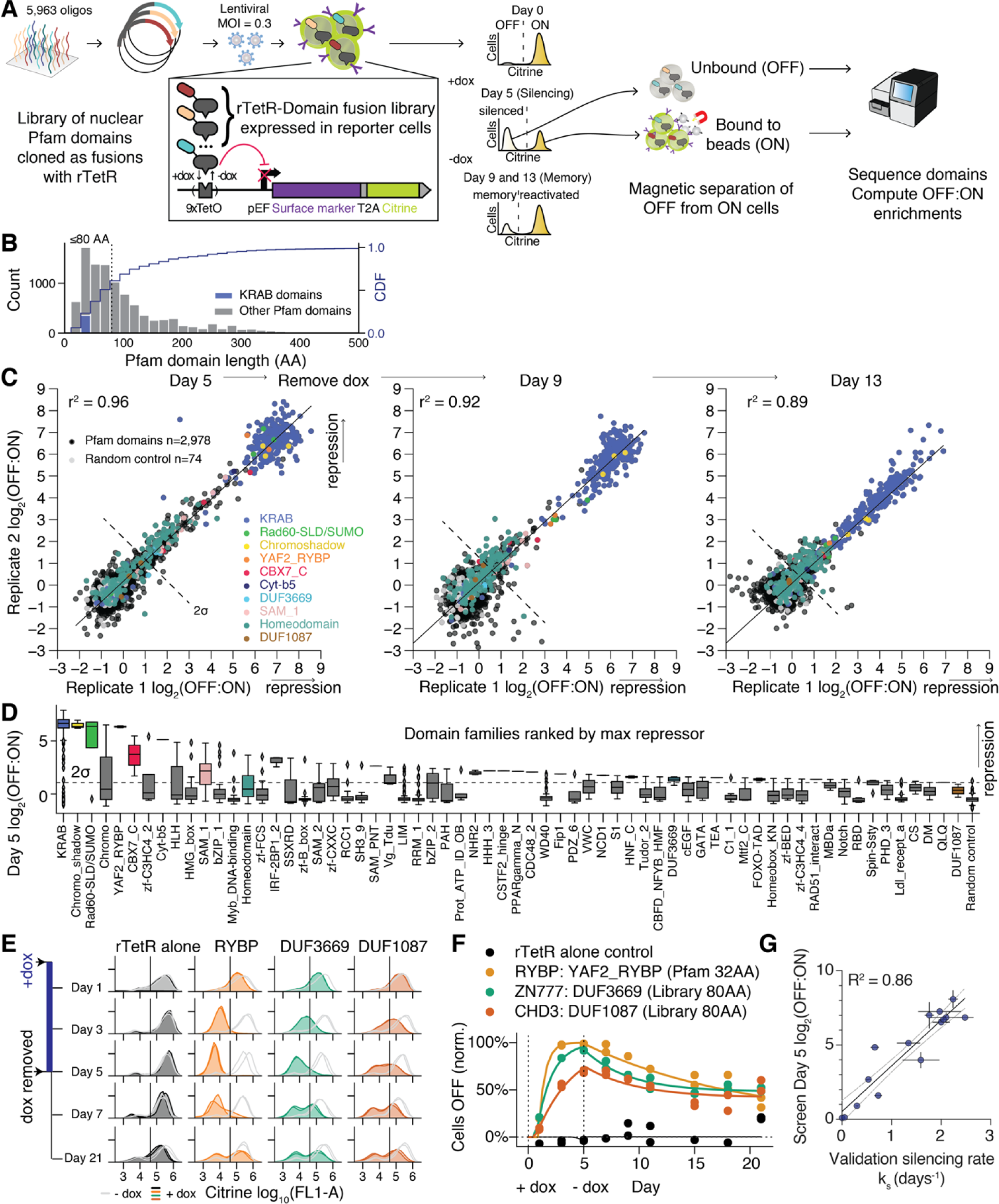
HT-recruit discovers hundreds of repressors in a screen of thousands of Pfam domains. A. Schematic of high-throughput recruitment assay (HT-recruit). A pooled library of Pfam domains is synthesized as 300-mer oligonucleotides, cloned downstream of the doxycycline (dox) -inducible rTetR DNA-binding domain, and delivered to reporter cells at a low multiplicity of infection (MOI) such that the majority of cells express a single rTetR-domain fusion. The repression reporter (inset) uses a strong pEF promoter that can be silenced by doxycycline-mediated recruitment of repressor domains via rTetR at the TetO sites. The reporter includes two components separated by a T2A self-cleaving peptide: a fluorescent citrine marker and an engineered synthetic surface marker (Igκ-hIgG1-Fc-PDGFRβ) that enables magnetic separation of ON from OFF cells. The reporter was stably integrated in the *AAVS1* safe harbor locus by homology-directed repair. The cells were treated with doxycycline for 5 days, ON and OFF cells were magnetically separated, and the domains were sequenced. Doxycycline was removed and additional time points were taken on Days 9 and 13 to measure epigenetic memory. B. Histogram of lengths of Pfam domains in human nuclear proteins. Domains ≤80 AA (dashed line) were selected for the library. Cumulative Distribution Function (CDF) is shown on the right-side axis. The KRAB family is shown as an example of an effector domain family. C. The reproducibility of log_2_(OFF:ON) ratios from two independently transduced biological replicates is shown and selected domain families are colored. The hit threshold at two standard deviations above the mean of the poorly expressed negative controls is shown with a dashed line. D. Boxplots of top repressor domain families, ranked by the maximum repressor strength at day 5 of any domain within the family. Line shows the median, whiskers extend beyond the high- and low-quartile by 1.5 times the interquartile range, and outliers are shown with diamonds. Dashed line shows the hit threshold. Boxes colored as in (D) for domain families highlighted in the text. E. Individual validations for RYBP domain and two Domains of Unknown Function (DUF) with repressor activity, measured by flow cytometry. Untreated cell distributions are shown in light grey and doxycycline-treated cells are shown in colors, with two independently-transduced biological replicates in each condition. The vertical line shows the citrine gate used to determine the fraction of cells OFF. F. Validation time courses fit with the gene silencing model: exponential silencing with rate k_s_, followed by exponential reactivation (**Methods**). Doxycycline (1000 ng/ml) was added on day 0 and removed on day 5 (N = 2 biological replicates). The fraction of mCherry positive cells with the citrine reporter OFF was determined by flow cytometry, as in (E), and normalized for background silencing using the untreated, time-matched controls. G. Correlation of high-throughput measurements at day 5 with the silencing rate k_s_ (R^2^ = 0.86, n = 15 domains, N = 2-3 biological replicates). Horizontal error bars are the standard deviation for the fitted rates, vertical error bars are the range of screen biological replicates, and dashed lines are the 95% confidence interval of the linear regression.

We next set out to curate a list of candidate transcriptional effector domains. First, we pulled sequences from the UniProt database for Pfam-annotated domains in human proteins that can localize to the nucleus (including non-exclusively nuclear-localized proteins). In total, we retrieved 14,657 domains. Of these, 72% were less than or equal to 80 amino acids (AA) long (**Figure 1B**), which made them compatible with pooled synthesis as 300 base oligonucleotides. For domains shorter than 80 AA, we extended the domain sequence on both ends with the adjacent residues from the native protein sequence in order to reach a length of 80 AA and avoid PCR amplification biases. We added 861 negative controls that were either random 80 AA sequences or 80 AA sequences tiled along the DMD protein with a 10 AA tiling window (**Materials and Methods**). The DMD protein is not localized in the nucleus (Chevron et al., 1994), and thus unlikely to feature domains with transcriptional activity. The library (**Table S1**) was cloned for lentiviral expression as a fusion protein with either the rTetR doxycycline-inducible DNA-binding domain alone, or with a 3X-FLAG-tagged rTetR (**Figure S2A and Table S2**) and delivered to K562 reporter cells (**Figure 1A**).

Before assaying for transcriptional activity, we determined which protein domains were well-expressed in K562 cells using a high-throughput approach (**Figure S2A**). We stained the library of cells with an anti-FLAG fluorescent-labeled antibody, sorted the cells into two bins (**Figure S2B**), extracted genomic DNA, and counted the frequency of each domain by amplicon sequencing (**Table S3**). We used the sequencing counts to compute the enrichment ratio in the FLAG_high_ versus FLAG_low_ population for each domain, as a measure of expression level. These measurements were reproducible between separately transduced biological replicates (r^2^ = 0.82, **Figure S2C**), and highly correlated with individual domain fusion expression levels measured by Western blot (r^2^=0.92, **Figures S2D and S2E**). As expected, native Pfam domains were significantly better-expressed than the random sequence controls (p<1e-5, Mann Whitney test), while the Pfam domains and the DMD tiling controls were similarly well-expressed (**Figure S2F**). We set a threshold to identify well-expressed domains with a FLAG_high_:FLAG_low_ ratio one standard deviation above the median of the random controls. By this definition, 66% of the Pfam domains were well-expressed; these domains were the focus of further analysis.

Next, we screened the Pfam domain library for transcriptional repressors. We treated the pooled library of cells with doxycycline for 5 days, which gives sufficient time after transcriptional silencing for the reporter mRNA and protein to degrade and dilute out due to cell division, resulting in a clear bimodal mixture of ‘ON’ and ‘OFF’ cells (**Figure S3A**). Then, we performed magnetic cell separation (**Figure S3A**) and domain sequencing, then computed the log_2_(OFF:ON) ratio for each library member using the read counts in the unbound and bead-bound populations (**Figure 1A**). For clarity, we refer to the bead-bound population as ‘ON’ and the unbound population as ‘OFF’. The measurements were highly reproducible between separately transduced biological replicates (r^2^ = 0.96, **Figure 1C**). We called domains as hits when they caused repression that was more than 2 standard deviations above the mean of the poorly expressed negative controls. This resulted in 446 repressor hits at day 5, with domains from 63 domain families (**Figure 1D**). These repressor domains are found in 451 human proteins, because in some cases the exact same domain sequence occurs in multiple genes. Known repressor domains (e.g. KRAB from human ZNF10, Chromoshadow from CBX5) from 10 domain families described as repressors or co-repressor-binding domains by Pfam were among the hits (**Table S4**), as expected.

To measure epigenetic memory, we took additional time points at days 9 and 13. The set of proteins containing hits was significantly enriched for transcription factors and chromatin regulators when compared to all nuclear proteins used in our library, but different categories of proteins were differentially enriched when classified by their memory levels (**Figure S3B**). Specifically, the repressors with high memory (cells remaining OFF) at day 13 were most enriched for C2H2 zinc finger transcription factors which include KRAB ZNF proteins, and the repressors with low memory were most enriched for homeodomain transcription factors which include the Hox proteins. Overall, the very high reproducibility and identification of expected positive control repressor domains among the hits suggested our screening method, which we called HT-recruit, yielded reliable results.

One of the strongest hits was the YAF2_RYBP, a domain present in the RING1- and YY1-binding protein (RYBP) and its paralog YY1-associated Factor 2 (YAF2), which are both components of the polycomb repressive complex 1 (PRC1) (Chittock et al., 2017; García et al., 1999). We individually tested the domain from the RYBP protein as annotated by Pfam (which is just 32 amino acids, thus shorter than the version we had synthesized in our 80 AA domain library) and confirmed rapid silencing of the reporter gene (**Figure 1E**). RYBP-mediated silencing was also demonstrated in a recent report of full-length RYBP protein recruitment in mouse embryonic stem cells (Moussa et al., 2019; Zhao et al., 2020). Our result establishes that the 32 AA RYBP domain, which has been shown by surface plasmon resonance to be the minimal required domain to bind the polycomb histone modifier enzyme RING1B (Wang et al., 2010), is sufficient to mediate silencing in cells.

To quantify repression kinetics, we gated the citrine level distributions to calculate a percentage of silenced cells, normalized the uniform low level of background silencing in the untreated cells, and then fit the data to a model with an exponential silencing rate during doxycycline treatment and an exponential decay (or reactivation) after doxycycline removal that plateaus at a constant irreversibly silent percentage of cells (**Figure 1F**). Using this approach, we also validated the repressor function of SUMO3, the Chromo domain from MPP8, the Chromoshadow domain from CBX1, and the SAM_1/SPM domain from SCMH1 (**Figures S3C - S3F**), which all had previous support for repressor function from recruitment or co-repressor binding assays (Chang et al., 2011; Chupreta et al., 2005; Frey et al., 2016; Lechner et al., 2000). Silencing rates from all individual measurements (for the repressor hits above and the other hits discussed below, **Figures S3C - S3K**) correlated well with the high-throughput measurements of silencing at day 5 (R^2^ = 0.86, **Figure 1G**). We performed these individual validations using a new variant of the DNA binding domain rTetR (SE-G72P) that was engineered to mitigate leakiness in the absence of doxycycline in yeast (Roney et al., 2016), and which we found is not leaky in human cells (**Figures S4A and S4B**), making it a useful tool for mammalian synthetic biology. This new rTetR variant has the same silencing strength at maximum doxycycline recruitment as the original rTetR (**Figure S4C**), which is also evidenced by the high correlation between individual validations and screen scores (**Figure 1G**). Together, these validation experiments demonstrated that HT-recruit both successfully identified bona fide repressors and quantified the repression strength for each domain with accuracy comparable to individual flow cytometry experiments.

### Identification of domains of unknown function that repress transcription

Over 22% of the Pfam domain families are labeled as Domains of Unknown Function (DUFs), while others are not named using this label but are nevertheless DUFs (El-Gebali et al., 2019). These domains have recognizable sequence conservation but lack experimental characterization. As such, our high-throughput domain screen offered the opportunity to associate initial functions with DUFs. First, we identified DUF3669 domains as repressor hits and individually validated one by flow cytometry (**Figures 1D - 1F**). These DUFs are natively found in KRAB zinc finger proteins, which is a gene family containing many repressive transcription factors. Concordant results demonstrating transcriptional repression after recruitment of two DUF3669 family domains were recently published (Al Chiblak et al., 2019), and our high-throughput results expand this finding to include the four remaining un-tested DUF3669 sequences (**Table S4**). The HNF3 C-terminal domain, HNF_C, is another DUF, although it has a more specific name because it is only found in Hepatocyte Nuclear Factors 3 alpha and beta (also known as FOXA1 and 2). We found the HNF_C domains from both FOXA1 and 2 were repressor hits (**Table S4**). Suggestively, they both include a EH1 (engrailed homology 1) motif, characterized by the FxIxxIL sequence, that has been nominated as a candidate repressor motif (Copley, 2005).

All three of the IRF-2BP1_2 N-terminal zinc finger domains (Childs and Goodbourn, 2003), an uncharacterized domain found in the interferon regulatory factor 2 (IRF2) co-repressors IRF2BP1, IRF2BP2, and IRF2BPL, were repressor hits (**Table S4**). The Cyt-b5 domain in the DNA repair factor HERC2 E3 ligase (Mifsud and Bateman, 2002) is another functionally uncharacterized domain that we validated as a strong repressor hit (**Figure S3G**). The SH3_9 domain in BIN1 is a largely uncharacterized variant of the SH3 protein-binding domain, which we also validated as a repressor (**Figure S3H**). BIN1 is a Myc-interacting protein and tumor suppressor (Elliott et al., 1999) that is also associated with Alzheimer’s disease risk (Nott et al., 2019). Concordant with our results, both full-length BIN1 and a Myc-binding domain deletion mutant were previously shown to repress transcription in a Gal4 recruitment assay in HeLa cells (Elliott et al., 1999), and the BIN1 yeast homolog hob1 has been linked to transcriptional repression and histone methylation (Ramalingam and Prendergast, 2007). In addition, we validated the repressor activity of the HMG_box domain from the transcription factor TOX and of the zf-C3HC4_2 RING finger domain from the polycomb component PCGF2 (**Figures S3I and S3J**). Lastly, DUF1087 is found in CHD chromatin remodelers and, although its high-throughput measurement was just below the screen significance threshold (**Figure 1D)**, we validated the CHD3 DUF1087 as a weak repressor by individual flow cytometry (**Figures 1E and 1F**). Together, these results demonstrate that high-throughput protein domain screens can assign initial functions to DUFs and expand our understanding of the functions of incompletely characterized domains.

### A random sequence with strong repressor activity

To our knowledge, random sequences have not previously been tested for repressor activity. We were surprised to find that one of the random 80 AA sequences, which were designed as negative controls, was a strong repressor hit with an average log_2_(OFF:ON) = 4.0, despite having a weak expression level below the threshold. Individual validation by flow cytometry confirmed that this sequence fully silenced the population of reporter cells after 5 days of recruitment with moderate epigenetic memory up to two weeks after doxycycline removal (**Figure S3K**). One additional random sequence showed a repression score marginally above the hit threshold and we did not attempt to validate it.

### Repressor KRAB domains are found in younger proteins

Our data provided an opportunity to analyze the function of all effector domains in the largest family of transcription factors: the KRAB domains. The KRAB gene family includes some of the strongest known repressor domains (such as the KRAB in ZNF10). Previous studies of a subset of repressive KRAB domains revealed that they can repress transcription by interacting with the co-repressor KAP1, which in turn interacts with chromatin regulators such as SETDB1 and HP1 (Cheng et al., 2014). However, it remains unclear how many of the KRAB domains are repressors, and whether the recruitment of KAP1 is necessary or sufficient for repression across all KRABs.

Our library included 335 human KRAB domains, and we found that 92.1% were repressor hits after filtering for domains that were well-expressed. We individually validated 9 repressor hit and 2 non-hit KRAB domains by flow cytometry and confirmed these categorizations in every case (**Figure S4D**). Then, we compared our domain recruitment results with previously published immunoprecipitation mass spectrometry data generated from full-length KRAB protein pulldowns (Helleboid et al., 2019) and found that all but one of the non-repressive KRABs were in proteins that do not interact with KAP1 (the one exceptional KRAB was lowly expressed in our study), and all of the repressor hit KRAB domains were KAP1 interactors (p<1e-9, Fisher’s exact test, **Figure 2A**). Furthermore, we analyzed available ChiP-seq and ChIP-exo datasets (ENCODE Project Consortium et al., 2020; Imbeault et al., 2017; Najafabadi et al., 2015; Schmitges et al., 2016) (**Table S5**) and found that repressive KRAB domains were from KRAB Zinc Finger proteins that co-localize with KAP1, in contrast to non-repressive KRAB domains (**Figure 2B**).

**Figure 2.**
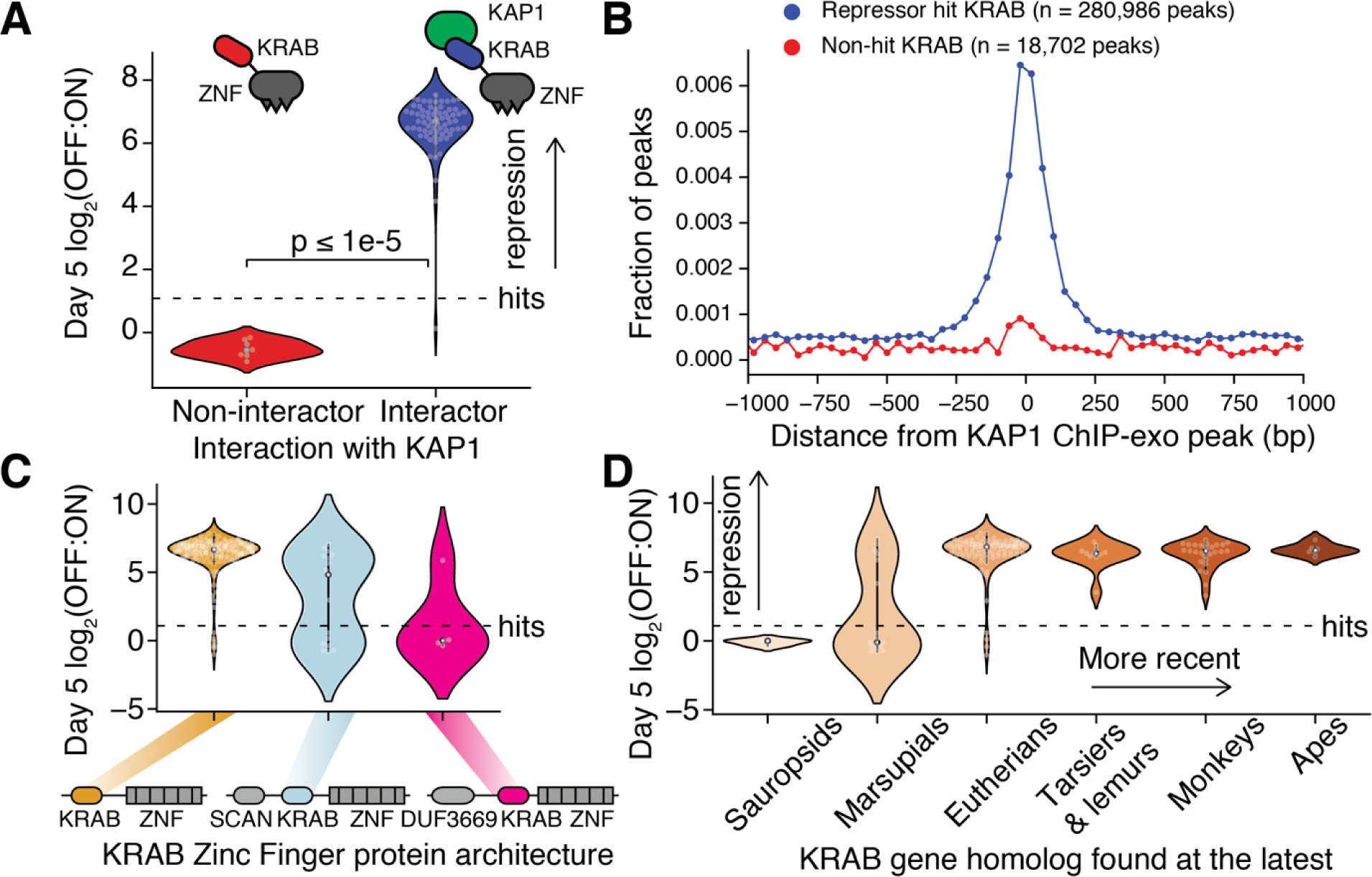
Repressor KRAB domains are found in younger KRAB-Zinc finger proteins that co-localize and bind to the KAP1 co-repressor. A. KRAB domain repression strength distributions (OFF:ON ratio after 5 days of recruitment) categorized by whether their KRAB Zinc Finger protein interacts significantly with co-repressor KAP1 by co-immunoprecipitation mass-spectrometry. Mass spectrometry dataset retrieved from (Helleboid et al., 2019). Each dot is a KRAB domain and the dashed line shows the hit threshold (N = 76 domains). B. Aggregate distance of ChIP peak locations of KRAB Zinc Finger proteins away from the nearest peaks of the co-repressor KRAB-associated Protein 1 (KAP1). KRAB domains are colored and classified by their status as hits or non-hits in the recruitment repression assay. Each dot shows the fraction of peaks in a 40 basepair bin. ChIP-seq and ChiP-exo data retrieved from (ENCODE Project Consortium et al., 2020; Imbeault et al., 2017; Najafabadi et al., 2015; Schmitges et al., 2016) (N = 150 hit KZFP ChIP datasets, N = 11 non-hit KZFP ChIP datasets). Only peaks where a single KRAB Zinc Finger binds are included (**Methods**). C. Repression measurements for KRAB domains that are natively found in KRAB Zinc Finger proteins with three different architectures. Each dot is a KRAB domain and the dashed line shows the hit threshold. D. Repression strength for KRAB domains from zinc finger genes of varying evolutionary age as determined by the most recent human ancestor with a genetic homolog (ages as reported in (Imbeault et al., 2017)). Each dot is a KRAB domain and the dashed line shows the hit threshold.

Interestingly, repressive KRAB domains were mostly found in proteins with the simplest domain architecture consisting of just a KRAB domain and a zinc-finger array, while the non-repressive KRAB domains were mostly found in genes that also include a DUF3669 or SCAN domain (**Figure 2C**). In fact, only one KRAB in a DUF3669-containing gene, ZNF783, was a repressor. ZNF783 is an uncharacterized DUF3669-KRAB-containing gene that uniquely lacks a zinc finger array (despite its name), suggesting it is distinctive among this class of transcription factors in both its effector function and its mode of localizing to targets.

The compound domain architecture that includes a SCAN or DUF3669 is more common in evolutionary old KRAB genes (Imbeault et al., 2017). Here, we observed a clear relationship between the evolutionary age of the KRAB genes and the KRAB repressor strength, with KRAB domains from genes pre-dating the marsupial-human common ancestor having no repressor activity, and KRAB domains from genes that evolved later consistently functioning as strong repressors (**Figure 2D**). Together, these results support a model of an ancient generation of non-repressor KRAB genes followed by a more recent massive expansion of repressor KRAB genes that recruit KAP1 to silence genomic targets.

### Deep mutational scan of the CRISPRi ZNF10 KRAB effector identifies mutations that modulate gene silencing

The KRAB domain from ZNF10 has been extensively used in synthetic biology applications for gene repression and is fused to dCas9 in the programmable epigenetic and transcriptional control tool known as CRISPR interference (Gilbert et al., 2014). To better understand its sequence-function relationships, we performed a deep mutational scan (DMS) of this KRAB domain using HT-recruit. We designed a library with all possible single substitutions and all consecutive double and triple substitutions (**Figure 3A**). To improve our ability to unambiguously align sequencing reads, we took advantage of variable codon usage to implement silent barcodes in the domain coding sequence such that the DNA sequences were more unique than the amino acid sequences (**Figure 3A**). We performed HT-recruit using the reporter and workflow in **Figure 1A**: 5 days of doxycycline induction and magnetic separation of ON and OFF cells at days 5, 9, and 13 (**Figure S5A**). These measurements were highly reproducible and showed a general trend of increasing deleteriousness with increasing mutation length from singles to triples, as expected (**Figure S5B**). Further, we compared these results with the KRAB amino acid conservation and found a striking correlation between conservation and deleteriousness of mutations (**Figure 3B**, bottom).

**Figure 3.**
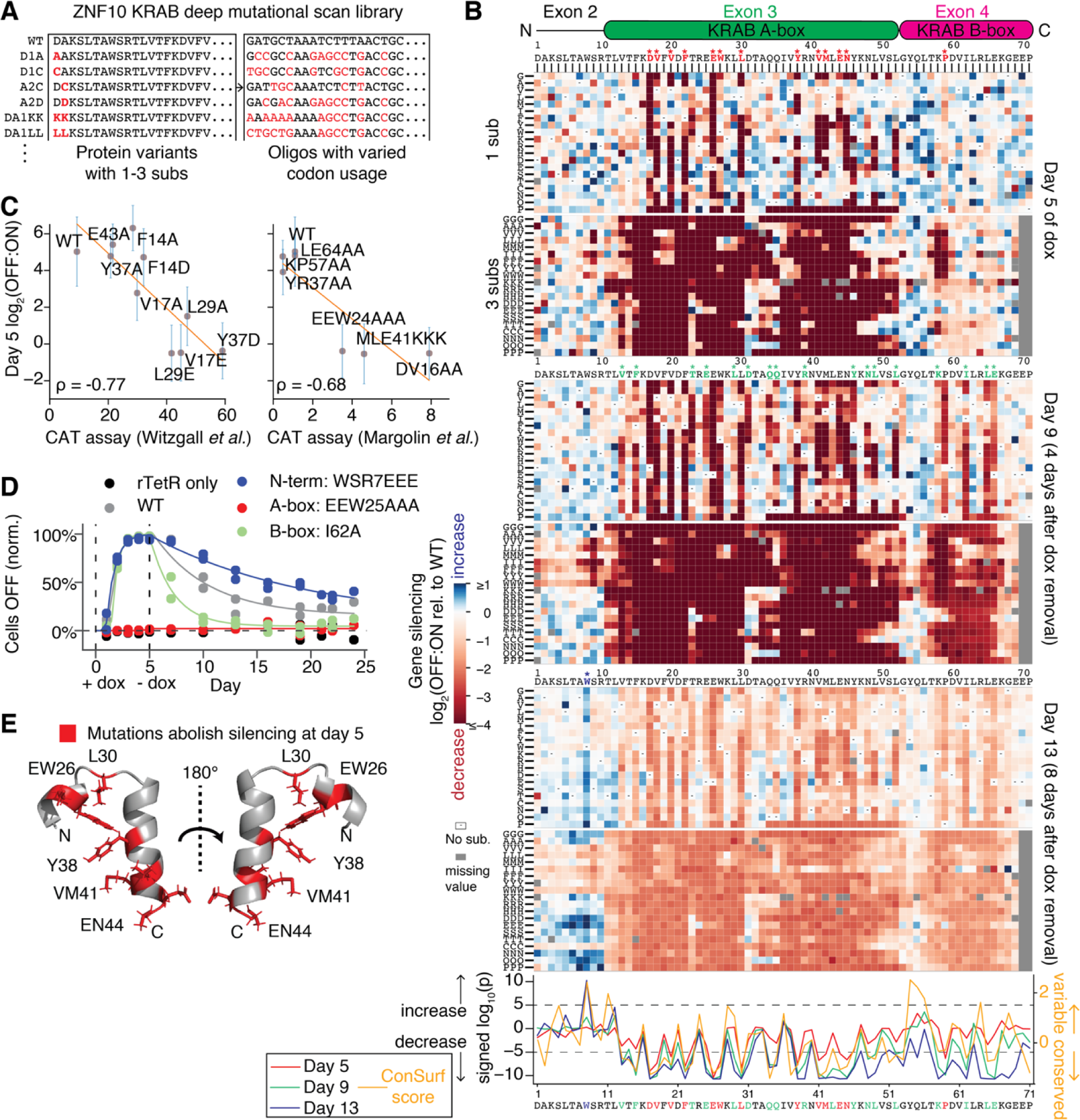
Deep mutational scan of the ZNF10 KRAB domain identifies substitutions that reduce or enhance repressor function. A. Design of a deep mutational scanning library including all single and consecutive double and triple substitutions in the KRAB domain of ZNF10. Red residues differ from the WT sequence shown on top. The DNA oligos are designed to be more distinct than the protein sequences by varying codon usage. B. (Top) All single and triple substitution (sub) variant repressor measurements relative to the WT are shown underneath a schematic of the KRAB domain. The N-terminal extension is encoded on exon 2, the KRAB A-box is encoded on exon 3, and the KRAB B-box is encoded on exon 4. Triple substitutions start at the position indicated. Dashes in a square indicate that it is the WT residue and grey indicates a missing value. Asterisks show significant residues, as defined below. (Bottom) For each position at each timepoint, the distribution of all single substitutions was compared to the distribution of wild-type effects (Wilcoxon rank sum test). Positions with signed log_10_(p)<-5 at day 5 are colored in red (highly significantly decrease in silencing), with signed log_10_(p)<-5 at day 9 but not day 5 are colored in green, and the position W8 with log_10_(p)>5 at day 13 is colored in blue (highly significant increase). Dashed horizontal lines show the hit thresholds. The sequence conservation ConSurf score is shown in orange. C. HT-recruit measurements correlate with previously published low-throughput recruitment CAT assay (Margolin et al., 1994; Witzgall et al., 1994). Vertical bars show the standard error from two biological replicates. A lower CAT assay value reflects a higher KRAB silencing activity. D. Individual time-courses of indicated rTetR-KRAB mutant fusions validate the effects of substitutions in the A/B-boxes and N-terminus. 1000 ng/ml doxycycline was added on day 0 and removed on day 5, the percentage of cells OFF was measured by flow cytometry, normalized for background silencing, and fit with the gene silencing model (**Methods**, N = 2 biological replicates of lentiviral infection). E. Residues that abolish silencing at day 5 when mutated are mapped onto the ordered region of the NMR structure of mouse KRAB A-box (PDB: 1v65).

The ZNF10 KRAB effector has 3 components: the A-box which is necessary for binding KAP1 (Peng et al., 2009), the B-box which is thought to potentiate KAP1 binding (Peng et al., 2007), and an N-terminal extension that is natively found on a separate exon upstream of the KRAB domain (**Figure 3B**). We found strong evidence for the necessity of the A-box, in which mutations at numerous positions dramatically lowered repressor activity relative to the wildtype sequence (**Figure 3B**). Several of these mutations had previously been tested with a recruitment CAT assay in COS and 3T3 cells; those data correlated well with measurements from our deep mutational scan in K562 cells (**Figure 3C**). We also individually validated the complete lack of silencing function in an A-box KRAB mutant (**Figure 3D**). The mutational impacts across the A-box appeared to be periodic, suggesting the angle of these residues along an alpha helix could be functionally relevant (**Figure 3B**). We called residues as necessary for silencing (p<1e-5, Wilcoxon rank sum test comparing distribution of all substitutions against wild-type at day 5) and found 12 necessary residues with strong mutational impacts in the A-box and one residue with significant but weak effects in the B-box (**Figure 3B**).

We mapped these substitutions onto an aligned mouse KRAB A-box structure (PDB: 1v65, 55% identity, 69% similarity in A-box [V13-Y54], **Figure S5C**) and found the necessary residues were similarly oriented in 3D space, suggestive of a binding interface (**Figures 3E and S5D**, red). We hypothesized that these residues were important for KAP1 binding. In agreement with this hypothesis, 10 out of 12 of these A-box residues were in fact shown to be necessary for KAP1 binding in a previous recombinant protein binding assay (Peng et al., 2009) using KRAB-O, which aligns to ZNF10 KRAB 12-71 (50% identity, 75% similarity) in a region containing all 12 of the necessary residues (red KRAB-O residues, **Figure S5C**). The remaining 8 out of 8 residues previously found unnecessary for binding were also not necessary for repression in our DMS (p<1e-4, Fisher’s exact test, grey KRAB-O residues, **Figure S5C**). We also inspected the DMS day 5 silencing scores for the individual single, double, and triple alanine substitutions used in the binding assay, and found perfect agreement: mutations that ablated binding also abolished silencing (Z-score<-4 compared to wild-type distribution), and mutations that did not affect binding also did not affect silencing (|Z-score|<0.6) (p<0.01, n = 12 mutations, Fisher’s exact test). This high validation rate, and their positioning in the 3D structure, suggests the remaining 2 out of 12 necessary A-box residues from the DMS (V41 and N45) could also be involved in KAP1 binding.

In contrast to the A-box, we found B-box mutations showed relatively little effect at the end of recruitment (day 5), with only one statistically significant position (P59) showing consistent but weak effects. Meanwhile P59 and 4 other positions (K58, I62, L65, E66) showed a significant effect on memory after doxycycline removal as measured at day 9 (**Figure 3B**). We performed individual validations for 4 significant positions and found that, as in the high-throughput experiment, the B-box mutants were strong gene silencers after day 5 of recruitment but showed reduced memory after doxycycline release (**Figures 3D and S5E**). To interpret this result, we considered our previously proposed gene silencing model in which silenced cells pass through a ‘reversibly silent’ state before entering an ‘irreversibly silent’ state (Bintu et al., 2016). We hypothesized that the B-box mutant memory reduction is the result of a moderate silencing speed reduction, resulting in fewer cells committing to the irreversibly silent state by day 5, and that the mutational impact on silencing speed was masked because reversibly silent and irreversibly silent cells are indistinguishable at day 5. To test this possibility, we repeated the silencing time course with a 100-fold lower dose of doxycycline in order to tune down the recruitment strength. In this regime, we observed that the B-box mutations reduced silencing speed before day 5 (**Figure S5E**). This result shows the B-box has a partial contribution to KRAB silencing speed.

Lastly, we found that only the KRAB N-terminus contained residues where many substitutions consistently enhanced silencing relative to wild-type (**Figure 3B**, blue, day 13 panels). In particular, nearly all substitutions for the tryptophan at position 8 led to higher numbers of cells silenced relative to wild-type at day 13 (which is the time point with the most dynamic range to detect silencing levels above wild-type). This was the only significant position for enhanced silencing (**Figure 3B**). We individually validated the memory enhancement for two of the highest-ranked of these mutants (WSR8EEE and AW7EE) with high-doxycycline recruitment (**Figures 3D and S5E**).

We hypothesized that this silencing enhancement could be the result of enhanced KRAB protein expression level. To investigate the relationship between protein expression level and KRAB silencing strength, we inspected our high-throughput FLAG-tag expression level measurements for the set of KAP1-binding KRAB domains and found that there was a significant correlation between KRAB expression level and silencing at day 13 (r^2^ = 0.49, **Figure S5F**). Most relevant to the deep mutational scan results, ZNF10 KRAB had lower expression levels compared to other KRAB domains that showed higher day 13 silencing levels, implying that it could be improved via mutations. Notably, the N-terminus is very poorly conserved (**Figure 3B**) and is in fact uniquely found in the KRAB from ZNF10 by BLAST, suggesting that stability-improving mutations in the N-terminus would be unlikely to interfere with KRAB function. In addition, across the entire domain expression dataset, we observed that higher tryptophan (W) frequency in a domain was negatively correlated with expression level while higher glutamic acid (E) frequency was positively correlated with expression level (**Figure S5G**). This amino acid composition trend further suggested that the N-terminal KRAB mutant enhancements could be due to improved expression level, as we specifically found that substituting out the tryptophan from KRAB position 8 enhanced its effector function and that this enhancement was most pronounced when substituting with glutamic acid. To test the hypothesis, we performed a Western blot for the ZNF10 KRAB variants and confirmed that the N-terminal glutamic acid substitution mutants were more highly expressed than the wild-type (**Figure S5H**). Together, these results demonstrated the use of a deep mutational scan both to map sequence-to-function for a human transcriptional repressor and to improve effectors by incorporating expression-enhancing substitutions into poorly conserved positions.

### Homeodomain repressor strength is colinear with Hox gene organization

The second largest domain family that included repressor hits in our screen was the homeodomain family. Homeodomains are composed of 3 helices and are sequence-specific DNA binding domains that make base contacts through Helix 3 (Lynch et al., 2006). In some cases, they are also known to act as repressors (Holland et al., 2007; Schnabel and Abate-Shen, 1996). Our library included the homeodomains from 216 human genes, and we observed that 26% were repressor hits. The repressors were found in 4 out of the 11 subclasses of homeodomains: PRD, NKL, HOXL, and LIM (**Figure 4A**). These recruitment assay results suggest that transcriptional repression could be a widespread, though not ubiquitous, function of homeodomain transcription factors.

**Figure 4.**
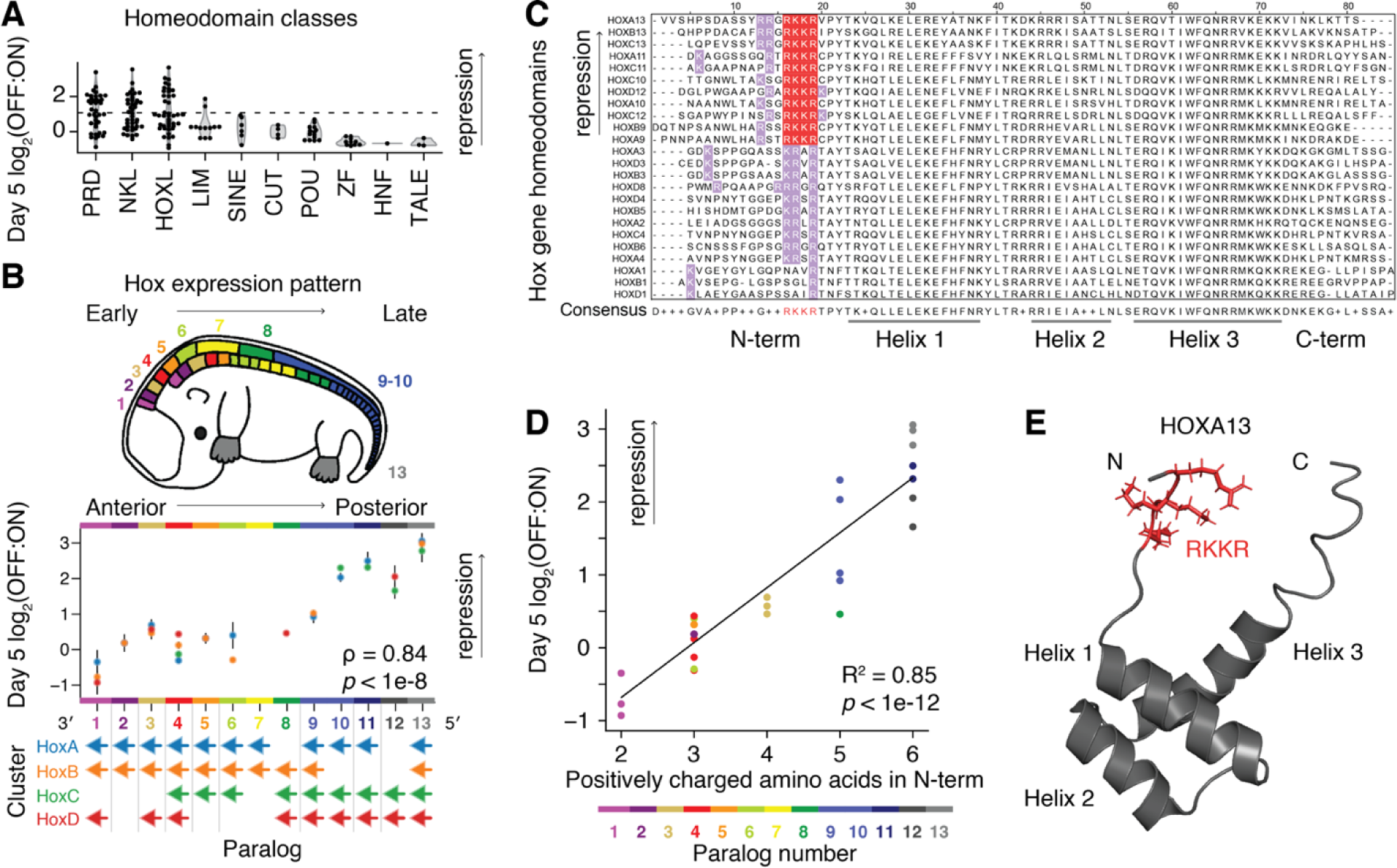
Hox homeodomain repression strength is colinear with Hox gene organization and associated with positive charge. A. Ranking of homeobox gene classes by median repression strength of their homeodomain at day 5. Horizontal line shows the hit threshold. None of the 5 homeodomains from the CERS class were well-expressed. B. Homeodomains from the *Hox* gene families. (Top) Hox gene expression pattern along the anterior-posterior axis is colored by Hox paralog number on an adapted embryo image (Hueber et al., 2010). Hox 11 and 12 are expressed both at the posterior end and along the proximal-distal axis of limbs (Wellik and Capecchi, 2003). (Middle) Repression strength after 5 days of dox. Dots are colored by the Hox cluster and the paralog number is colored as in the embryo diagram. Spearman’s rho and p-value were computed for the relationship between the paralog number and repressor strength across all Hox genes. (Bottom) Colored arrows represent the genes found in the four human Hox clusters and point in the direction of Hox gene transcription from 5′ to 3′. Grey bars separate gene sequence similarity groups as previously classified (Hueber et al., 2010). C. Multiple sequence alignment of Hox homeodomains, with stronger repressors at the top (as ranked by OFF:ON ratio at day 5), showing the RKKR motif highlighted in red. Other basic residues within the N-terminal arm are colored in lavender. D. Correlation between the number of positively charged residues in the N-terminal arm upstream of Helix 1 of each Hox homeodomain and the average repression at day 5. Dot color shows paralog number. E. NMR structure of the HOXA13 homeodomain retrieved from PDB ID: 2L7Z, with RKKR motif highlighted in red. The sequence from G15 to S81, using the coordinates from the multiple sequence alignment, is shown.

Next, we inspected the HOXL subclass results more closely. This subclass contains the *Hox* genes, a subset of 39 homeodomain transcription factors that are master regulators of cell fate and specify regions of the body plan along the anterior-posterior axis during embryogenesis. These genes are found in four *Hox* paralog clusters (A to D) that are arranged co-linearly from 3′ to 5′ corresponding to the temporal order and spatial patterning of their expression along the anterior-posterior axis (Gilbert, 1971). Interestingly, we observed that the repressor strength of their homeodomains was also collinear with their arrangement in the *Hox* clusters, such that the more 5′ gene homeodomains were stronger repressors (Spearman’s ρ = 0.82, **Figure 4B**). This correlation suggests a possible link between homeodomain repressor function and *Hox* gene expression timing and anterior-posterior axis spatial patterning.

We next sought to identify the sequence determinants of the Hox homeodomain repression strength gradient. Multiple sequence alignment of the Hox homeodomains revealed an RKKR motif present in the N-terminal arm of the 11 strongest repressor domains (**Figure 4C**). The motif resides in a basic context in the strongest repressors, while the lower ranked domains lack the motif but still contain some basic residues in the disordered N-terminal arm, resulting in a significant correlation between repression strength and the number of positively charged amino acids arginine and lysine (R^2^ = 0.85, **Figures 4C - 4E**).

We then investigated whether the RKKR motif and positive charge were associated with repression in other domains. Outside the Hox homeodomains, 99.5% of the repressor hits in the Pfam nuclear protein domain library do not contain the RKKR motif, while many non-hits do. Also, there was no correlation between net domain charge and repression strength at day 5 when considering the full library of domains (R^2^ = 0.04). Together, these results suggest the RKKR motif and charge contribute to Hox homeodomain repression in the recruitment assay, but they are not sufficient for repression when found in the context of other domains.

### Discovery of transcriptional activators by HT-recruit to a minimal promoter

Next, we set out to discover transcriptional activator domains using HT-recruit. We established a reporter K562 line with a weak minimal CMV (min CMV) promoter that could be activated upon recruitment of fusions between rTetR and activation domains (**Figure 5A**). To perform the activator screen, we used lentivirus to deliver the nuclear Pfam domain library to these reporter cells, induced rTetR-mediated recruitment with doxycycline for 48 hours, magnetically separated the cells (**Figure S6A**), and sequenced the domains in the two resulting cell populations. We computed an enrichment ratio from the sequencing counts in the bead-bound (ON) and unbound (OFF) populations for each domain as a measure of transcriptional activation strength and called hits that were two standard deviations past the mean of the poorly expressed negative controls (**Figure 5B**). The hits included three previously known transcriptional activation domain families that were present in our library: FOXO-TAD from FOXO1/3/6, LMSTEN from Myb/Myb-A, and TORC_C from CRTC1/2/3. Encouragingly, activation strength measurements for the hits were highly reproducible between separately transduced biological replicates (r^2^ = 0.89, **Figure 5B**). This second screen with the short nuclear domain library established that HT-recruit can be used to measure either activation or repression by changing the reporter’s promoter.

**Figure 5.**
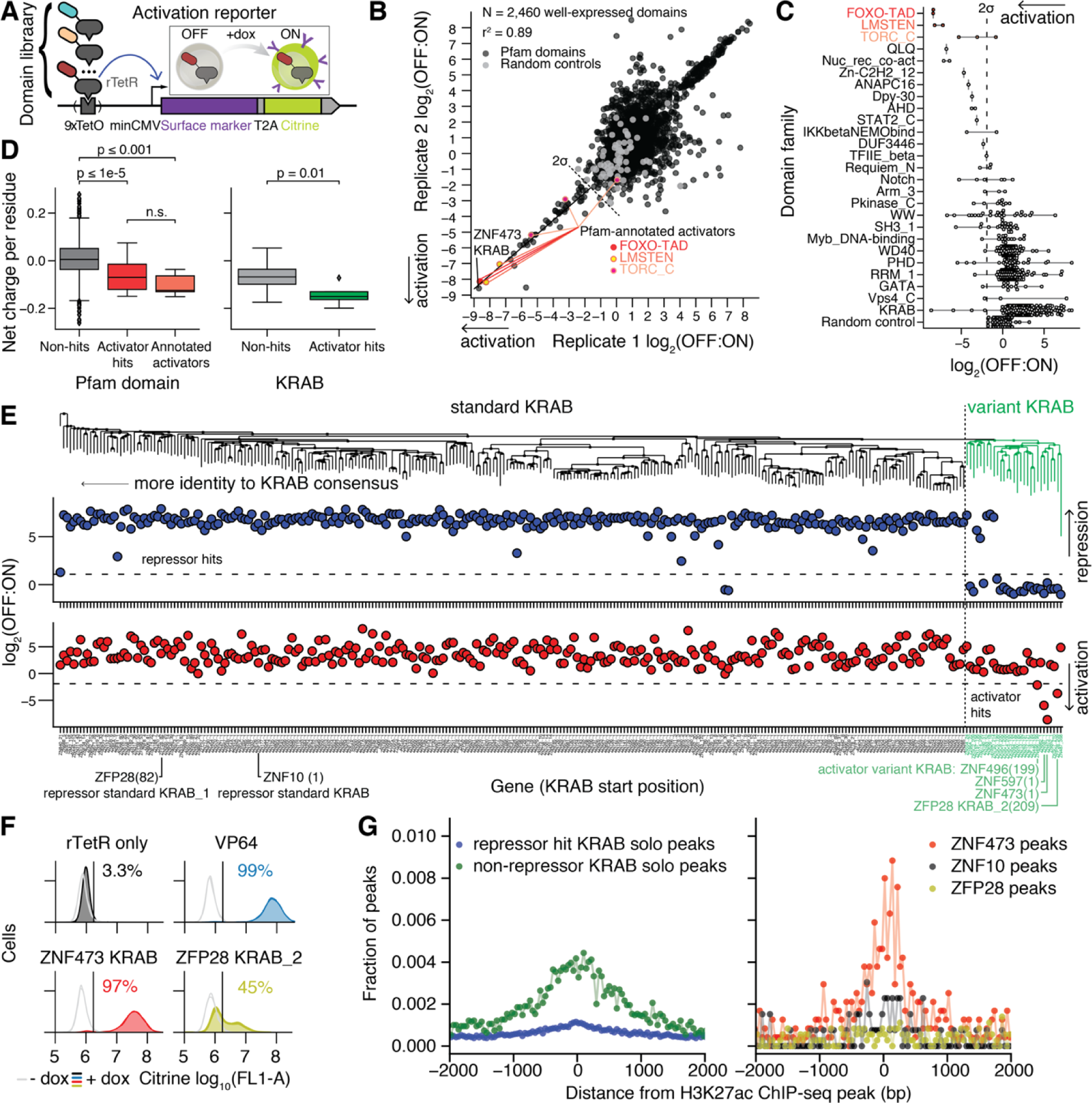
HT-recruit discovers activator domains. A. Schematic of the activation reporter which uses a weak minCMV promoter that can be activated by doxycycline-mediated recruitment of activating effector domains fused to rTetR. B. Reproducibility of high-throughput activator measurements from two independently transduced biological replicates. The pool of cells containing the activation reporter in (A) were transduced with the nuclear domain library and treated with doxycycline for 48 hours; ON and OFF cells were magnetically separated, and the domains were sequenced. The ratios of sequencing reads from the OFF vs. ON cells are shown for domains that were well-expressed. Pfam-annotated activator domain families (FOXO-TAD, Myb LMSTEN, TORC_C) are colored in shades of red. A line is drawn to the strongest hit, the KRAB domain from ZNF473. The hit threshold is a dashed line drawn two standard deviations below the mean of the poorly expressed domain distribution. C. Rank list of domain families with at least one activator hit. Families previously annotated as activators in Pfam are in red. The dashed line represents the hit threshold, as in (B). Only well-expressed domains are shown. D. Acidity of effector domains from the Pfam library, calculated as net charge per amino acid. (Left) Comparison of the non-hit, well-expressed Pfam domains (except KRAB and annotated activators) with the activator hits. The Pfam-annotated activator domain families are shown as a group as a positive control (orange). (Right) Comparison of the activator hits and non-hits from the KRAB domain family. P-values from Mann-Whitney test shown with bars between compared groups. n.s. = not significant (p>0.05). E. Phylogenetic tree of all well-expressed KRAB domains with the sequence-divergent variant KRAB cluster shown in green (top). High-throughput recruitment measurements for repression at Day 5 are shown in blue (middle) and measurements for activation are shown in red (bottom). Dashed horizontal lines show hit thresholds. An example repressor KRAB from ZNF10, the repressor KRAB_1 from ZFP28, and all of the activator KRAB domains are called out with larger labels. The KRAB domain start position is written in parentheses. F. Individual validation of variant KRAB activator domains. rTetR(SE-G72P)-domain fusions were delivered to K562 reporter cells by lentivirus and selected for with blasticidin, cells were treated with 1000 ng/ml doxycycline for 2 days, and then citrine reporter levels were measured by flow cytometry. Untreated cell distributions are shown in light grey and doxycycline-treated cells are shown in colors, with two independently-transduced biological replicates in each condition. The vertical line shows the citrine gate used to determine the fraction of cells ON and the average fraction ON for the doxycycline-treated cells is shown. G. Distance of ChIP peak locations of KRAB Zinc Finger proteins away from the nearest peaks of the active chromatin mark H3K27ac. KRAB proteins are classified by their status as hits (blue) or non-hits (green) in the repressor screen at day 5 (left). In addition, data is shown individually for ZNF10 which contains a repressor hit KRAB (black), ZNF473 which contains an activator hit KRAB (red), and ZFP28 which contains both an activator hit and a repressor hit KRAB (yellow) (right). Each dot shows the fraction of peaks in a 40 basepair bin. ChIP-seq and ChiP-exo data retrieved from (ENCODE Project Consortium et al., 2020; Imbeault et al., 2017; Najafabadi et al., 2015; Schmitges et al., 2016). Only solo peaks, where a single KRAB Zinc Finger binds, are included for the aggregated data (blue and green dots, left), but all peaks are included for the individual proteins because the number of solo peaks is low for each individual protein (red, black, and yellow dots, right) (**Methods**).

In total, we found 48 hits from 26 domain families (**Table S4**). Beyond the three known activator domain families above, the remaining families with an activator hit were not previously annotated on Pfam as activator domains (**Figure 5C**). Overall, we found fewer activators than repressors, which may simply be because activators are often disordered or low-complexity regions (Liu et al., 2006) that are frequently not annotated as Pfam domains. However, the proteins containing activator domains were significantly enriched for gene ontology terms such as ‘positive regulation of transcription’ and the strongest enrichment was for ‘signaling’, which reflects that many of their source proteins are activating factors (**Figure S6B**). Further, the hits were significantly more acidic than non-hits (p ≤ 1e-5, Mann Whitney test, **Figure 5D**), a common property in activation domains (Mitchell and Tjian, 1989; Staller et al., 2018).

Several hits were not sourced from sequence-specific transcription factors where classical activator domains are expected but were instead nonclassical activators from co-activator and transcriptional machinery proteins including Med9, TFIIEβ, and NCOA3. In particular, the Med9 domain, whose ortholog directly binds other mediator complex components in yeast (Takahashi et al., 2009), was a strong activator with an average log_2_(OFF:ON) = -5.5, despite its weak expression level (**Table S4**). Nonclassical activators have previously been reported to work individually in yeast (Gaudreau et al., 1999) but only weakly when individually recruited in mammalian cells (Nevado et al., 1999). One exception is TATA-binding protein (Dorris and Struhl, 2000). By screening more nonclassical sequences, we were able to find more exceptions to this notion.

We then set out to individually validate activator hits by flow cytometry. For all tested domains, we confirmed doxycycline-dependent activation of the reporter gene using both the extended 80 AA sequence from the library and the trimmed Pfam-annotated domain (**Figure S6C**). As expected, the previously annotated FOXO-TAD and LMSTEN were strong activators, in both their extended and trimmed versions. We also validated the activator function of DUF3446 from the transcription factor EGR3 and the largely uncharacterized QLQ domain from the SWI/SNF family SMARCA2 protein. Further, we confirmed that the Dpy-30 motif domain, a DUF found in the Dpy-30 protein, is a weak activator. Dpy-30 is a core subunit of histone methyltransferase complexes that write H3K4me3 (Hyun et al., 2017), a chromatin mark associated with transcriptionally active chromatin regions (Sims et al., 2003). In total, we tested 11 hit domains (including nonclassical hits Med9 and Nuc_rec_co-act from NCOA3) and found that all of them significantly activated the reporter, when using the extended 80 AA sequence from the library. Together, the screen and validations demonstrate that our unbiased nuclear protein domain library can be productively re-screened to uncover domains with distinct functions, and that a diverse set of domains beyond classical activation domains (and including DUFs) can activate transcription upon recruitment.

### Discovery of KRAB activator domains

Surprisingly, the strongest activator in the library was the KRAB domain from ZNF473 (**Figure 5B**). Three other KRAB domains (from ZFP28, ZNF496, and ZNF597) were also activator hits, all of which were stably expressed and not repressors. One of these domains, from ZNF496, had previously been reported as an activator when recruited individually in HT1080 cells (Losson and Nielsen, 2010). Interestingly, ZFP28 contains two KRAB domains; KRAB_1 is a repressor and KRAB_2 is an activator. Previous affinity-purification/mass-spectrometry performed on full-length ZFP28 identified significant interactions with both repressor and activator proteins (Schmitges et al., 2016). The activator KRAB domains are significantly more acidic than non-activator KRABs (p = 0.01, Mann Whitney test, **Figure 5D**). Sequence analysis showed they were divergent from the consensus KRAB sequence while sharing homology to each other and formed a variant KRAB subcluster (**Figure 5E**). Previous phylogenetic analysis has linked the variant KRAB cluster to a lack of KAP1 binding and older evolutionary age (Helleboid et al., 2019). More specifically, two of the activator KRAB source proteins (ZNF496 and ZNF597) have previously been tested with co-immunoprecipitation mass-spectrometry and were not found to interact with KAP1 (Helleboid et al., 2019).

We individually validated the KRAB from ZNF473 as a strong activator and KRAB_2 from ZFP28 as a moderate strength activator (**Figure 5F**), when using the same 80 AA sequence centered on the KRAB domains that was used in our library. Further, we found the trimmed 41 AA KRAB from ZNF473 was sufficient for strong activation, while the trimmed 37 AA KRAB_2 from ZFP28 did not activate, implying some of the surrounding sequence is required for activation (**Figure S6C**). Next, we inspected available ChiP-seq and ChIP-exo datasets (ENCODE Project Consortium et al., 2020; Imbeault et al., 2017; Najafabadi et al., 2015; Schmitges et al., 2016) (**Table S5**) and found that ZNF473 co-localizes with the active chromatin mark H3K27ac, in contrast to the repressive ZNF10 (**Figure 5G**). Upon manual inspection, we found that the most significant ZNF473 peaks were found near the transcription start site of genes (*CASC3, STAT6, WASF2, ZKSCAN2*) and a lncRNA (*LINC00431*). Meanwhile, ZFP28 does not co-localize with H3K27ac, perhaps indicating its KAP1-binding repressor KRAB_1 domain is generally the dominant effector over its moderate strength activator KRAB_2 domain. Looking beyond these individual KRAB proteins, we found the zinc finger proteins that contain a repressor KRAB do not co-localize with H3K27ac while the non-repressive KRAB proteins as a group do include co-localized peaks (**Figure 5G**). Together, our results support that variant KRAB proteins are functionally diverse, sometimes functioning as transcriptional activators.

### Tiling library uncovers effector domains in unannotated regions of nuclear proteins

Pfam annotations provided one useful means of filtering the nuclear proteome to generate a relatively compact library, but Pfam is likely currently missing many of the human effector domains. In order to discover effector domains in unannotated regions of proteins, we designed a tiling library by curating a list of 238 proteins from silencer complexes (**Table S1**) and tiling their sequences with 80 amino acids separated by a 10 amino acid tiling window (**Figure 6A**). We performed high-throughput recruitment to the strong pEF reporter and took time points after 5 days of doxycycline to measure silencing, and again at day 13 (8 days after doxycycline release) to measure epigenetic memory (**Figure S7A**). 4.3% of the tiles scored as hits at day 5 (**Figure 6B**) and their repressor strength measurements were reproducible (r^2^ = 0.72, **Figure S7B and Table S4**). Altogether, the tiling screen found short repressor domains in 141/238 proteins. Some of these hits include positive controls overlapping annotated domains: for example, by tiling ZNF57 and ZNF461, we can identify the KRAB domains of these transcription factors as repressive effectors, and not the rest of the sequence (**Figure S7C**). Similarly, the tiling strategy identified the RYBP repressive domain annotated by Pfam, and both the 80 AA tile and the 32 AA Pfam domain silenced with similar strength and epigenetic memory in individual validations (**Figure S7D**). We also identified and validated repressors in REST (overlapping the CoREST binding domain (Ballas et al., 2001)), DNMT3b (overlapping the DNMT1 and DNMT3a binding domain (Kim et al., 2002)), and CBX7 (overlapping the PcBox that recruits PRC1 (Li et al., 2010)) (**Figures S7E - S7G**). Another category of tiling hits is not annotated as domains in Pfam, but we found previous reports of their repressor function in the literature. For example, we found that amino acids 121-220 of CTCF have a strong repressive function in the screen and when validated individually (**Figures 6C and 6E**), consistent with previous recruitment studies in HeLa, HEK293, and COS-7 cells (Drueppel et al., 2004). Together, these results established that high-throughput recruitment of protein tiles is an effective strategy to identify bonafide repressor domains.

**Figure 6.**
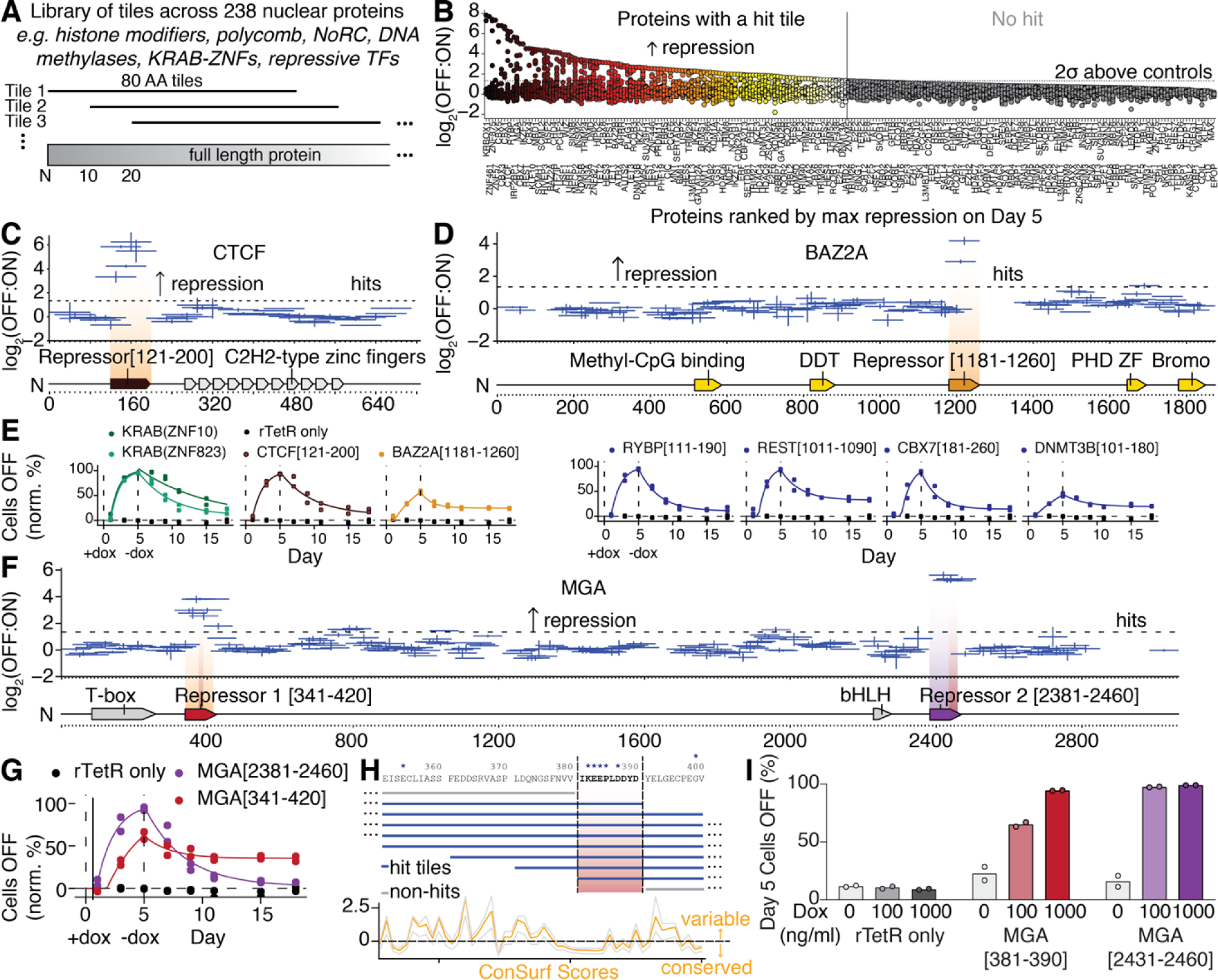
Tiling screen discovers compact repressor domains within nuclear proteins. A. Schematic of 80 AA tiling library covering a curated set of 238 nuclear-localized proteins. These tiles were fused to rTetR and recruited to the reporter, using the same workflow as in **Figure 1A** to measure repression strength. B. Tiled genes ranked by maximum repressor function at day 5 shown with a dot for each tile. Hits are tiles with a log_2_(OFF:ON) ≥ 2 standard deviations above the mean of the negative controls. Genes with a hit tile are colored in a gradient and genes without any hit tiles are colored in grey. C. Tiling CTCF. Diagram shows protein annotations retrieved from UniProt. Horizontal bars show the region spanned by each tile and vertical error bars show the standard error from two biological replicates of the screen. The strongest hit tile is highlighted with a vertical gradient and annotated as a repressor domain (orange). D. Tiling BAZ2A (also known as TIP5). E. Individual validations. Lentiviral rTetR(SE-G72P)-tile fusions were delivered to K562 reporter cells, cells were treated with 100 ng/ml doxycycline for 5 days (between dashed vertical lines), and then doxycycline was removed. Cells were analyzed by flow cytometry, the fraction of cells with citrine reporter OFF was determined and the data fit with the gene silencing model (**Methods**) (N = 2 biological replicates). Two KRAB repressor domains are shown as positive controls. The tiling screen data that corresponds to the validations shown on the right (blue curves) is shown in **Figure S7**. F. Tiling MGA. Two repressor domains are found outside the previously annotated regions and labeled as Repressor 1 and 2 (dark red, purple). The minimized repressor regions at the overlap of hit tiles are highlighted with narrow red vertical gradients. G. The maximal strength repressor tiles from two peaks in MGA were individually validated with the method described in (E) (N = 2 biological replicates). H. The MGA repressor 1 sequence was minimized by selecting the region shared in common between all hit tiles in the peak, shown between dashed vertical lines and shaded in red. The protein sequence conservation ConSurf score is shown below with an orange line and the confidence interval (the 25th and 75th percentiles of the inferred evolutionary rate distribution) is shown in grey. The asterisks mark residues that are predicted to be functional (highly conserved and exposed) by ConSurf. The repressor 2 sequence was minimized with the same approach and also overlaps a region with predicted functional residues (data not shown). I. The MGA effectors were minimized to 10 and 30 AA sub-tiles, as shown in (H), cloned as lentiviral rTetR(SE-G72P)-tile fusions, and were delivered to K562 reporter cells. After selection, cells were treated with 100 or 1000 ng/ml doxycycline for 5 days and the percentages of cells with the Citrine reporter silenced were measured by flow cytometry (N = 2 biological replicates).

Excitingly, we also discovered novel unannotated repressor domains. For example, BAZ2A (also known as TIP5) is a nuclear remodeling complex (NoRC) component that mediates transcriptional silencing of some rDNA (Guetg et al., 2010), but does not have any annotated effector domains. Our BAZ2A tiling data showed a peak of repressor function in a glutamine-rich region and we individually validated it as a moderate strength repressor (**Figures 6D and 6E**). We found repressor tiles in unannotated regions of three TET DNA demethylases (TET1/2/3) (**Table S4**). Unexpectedly, we also identified repressor tiles in the control protein DMD, which we validated by flow cytometry (**Figure S7H**).

Finally, we focused on the transcription factor MGA which is thought to repress transcription by binding the genome at E-box motifs and recruiting the non-canonical polycomb 1.6 complex (Blackledge et al., 2014; Jolma et al., 2013; Stielow et al., 2018); however, the domains responsible for silencing activity in this protein had not been identified. Our MGA tiling experiment revealed two domains with repressor function, located adjacent to the two known DNA binding domains, which we call here Repressor 1 and Repressor 2 (**Figure 6F**). We individually validated these repressor domains and observed distinct dynamics of silencing and degrees of memory; the first domain (amino acids 341-420) featured slow silencing but strong memory, while the second domain (amino acids 2381-2460), featured rapid silencing but weak memory with fast reactivation (**Figure 6G**). These are the first effector domains isolated from a protein in the ncPRC1.6 silencing complex, to our knowledge.

Next, we attempted to identify the minimal necessary sequence for repressor function in each independent domain by examining the overlap in all tiles covering a protein region that shows repressor function and determining which contiguous sequence of amino acids is present in all the repressive tiles (**Figure 6H**). Using this approach, we generated two candidate minimized effector domains for MGA: the 10 amino acid sequence MGA[381-390] and the 30 amino acid sequence MGA[2431-2460], which both overlapped conserved regions with ConSurf-predicted functional exposed residues. Individual validation experiments demonstrated that both minimized candidates can efficiently silence the reporter (**Figure 6I**).

## Discussion

The team behind the Pfam database has reported that “over a quarter of Pfam entries lack an experimentally validated function, highlighting the desperate need for more high-throughput functional screening of proteins” (El-Gebali et al., 2019). This need is particularly pressing in human cells, which can be more difficult than simpler model organisms like yeast for developing high-throughput assays, but which are also a necessary model for better understanding human biology and disease. Here we use HT-recruit to associate functions with Pfam domains including domains of unknown function (e.g. DUF1087, DUF3669, DUF3446, HNF_C, Dpy-30) and discover effectors in unannotated regions of proteins including BAZ2A and MGA. We find repressor domains in at least 552 human proteins and activators in 48 proteins. This resource of >600 effector domains could be used in the short term for interpreting the roles of proteins in the human nucleus based on the capacity of their domains to activate or repress transcription. In the longer term, these discoveries set the stage for deepening our knowledge of transcriptional effector mechanisms through detailed analysis of individual effectors and for enabling synthetic biology approaches to manipulate transcription.

### Functional divergence in the KRAB Zinc Finger Transcription Factor family

Our approach afforded us the opportunity to comprehensively assess the transcriptional regulatory function of the complete family of human KRAB domains. KRAB domains are generally thought of as repressors, and indeed we find that 92.1% of 335 KRAB domains are repressor hits. However, these results do not explain the long-term maintenance over millions of years of evolution of the ancient KRAB genes without repressor KRAB activity. The KRAB domains in this category are divergent in sequence, form a distinct cluster in a phylogeny of KRAB domains, are found in proteins that do not interact with KAP1, and have been called ‘variant KRABs’ (Helleboid 2019). One possibility is that these KRAB genes also act as repressors, although their KRAB domain alone is insufficient to mediate repression. Four of these non-repressor KRAB domains are in proteins that also contain a domain of unknown function, DUF3669. All four of the DUF3669 domains in those genes are repressor hits, albeit with lower strength than standard KRAB domains. On the other hand, we also found 4 variant KRAB domains (in ZNF473, ZFP28, ZNF496, and ZNF597) were activator hits. In further concordance with our data, bioinformatic investigation into the origins of the KRAB domain nominated transcriptional activation as the original KRAB function, due to ancestral KRAB sequence similarity with the Meisetz gene encoding for the PRDM9 activator (Birtle and Ponting, 2006). Together, these results support an evolutionary history in which the ancestral KRAB domain was non-repressive (and may have been activating), with repressive function originating around the time of the most recent human common ancestor with marsupials and subsequently becoming the predominant KRAB function among KRAB Zinc Finger proteins.

### Homeodomain repressors in *HOX* genes

We also gathered comprehensive measurements of homeodomain repressor strength and observed a significant collinear relationship between Hox gene cluster organization and repression strength. While it remains to be seen how this gradation of repressor strength affects *Hox* gene biological roles in settings such as embryogenesis, it is tempting to speculate that it could contribute to posterior prevalence. Briefly, posterior prevalence can be defined as the tendency for the more 5′ Hox gene to dominate the phenotype of a cell that expresses multiple *Hox* genes (Duboule and Morata, 1994). Experiments in fly and mouse have identified the homedomain as a mediator of posterior prevalence in some contexts, by performing homeodomain swaps from posterior to anterior *Hox* genes (Mann and Hogness, 1990; Zhao and Potter, 2001). One possibility suggested by our data is that the *Hox* genes repress one another, and the stronger repressor function of the 5′ *Hox* gene promotes its tendency to dominate the 3′ *Hox* gene. This would be compatible with the widespread evidence that expression of more 5′ *Hox* genes is correlated with reduced expression of more 3′ *Hox* genes (reviewed in (Mallo and Alonso, 2013)). Meanwhile, a number of experiments have demonstrated *Hox* regulatory mechanisms outside of the Hox proteins themselves, including miRNA, lncRNA and translational regulatory mechanisms (Mallo and Alonso, 2013). Further study is needed to elucidate if and how Hox homeodomain repression relates to these other mechanisms and contributes to posterior prevalence.

Hox homeodomain repression strength correlated with its N-terminal positive charge and the presence of an RKKR motif, which suggests a possible repression mechanism. Arginine residues in basic motifs are important for nucleolar localization (Birbach et al., 2004; Martin et al., 2015). Nucleophosmin mediates nucleolar integration of proteins containing these motifs via phase separation (Mitrea et al., 2016). While Hox homeoproteins have been shown to localize to the nucleolus before, the function of this localization was unclear (Corsetti et al., 1995). The nucleolus is highly enriched in components necessary for ribosomal synthesis, but excludes RNA polymerase II (Pol II) mRNA transcription (Sirri et al., 2008). Additionally, genomic regions in close proximity to the nucleolus are associated with repressive chromatin modifications (Bersaglieri and Santoro, 2019) Therefore, localization of a Hox homeodomain and its target gene to the nucleolus could serve as a mechanism for repression of Pol II-dependent genes such as our reporter. These observations provide a testable hypothesis for a tunable repression mechanism in Hox homeodomains whereby the incorporation of additional positive charges increases the repression strength of the more 5′ *Hox* genes.

### Deep mutational scanning domains in human cells

HT-recruit presents an approach to not only expand our catalog of human transcriptional effectors, but also to map their sequence-function relationships with mutational libraries. Deep mutational scans (DMS) are a relatively recent technology that has mostly been applied using yeast, bacteria, bacteriophage display (Fowler and Fields, 2014), and more recently demonstrated with both transient transfection (Heredia et al., 2018) and lentiviral delivery to human cells (Kotler et al., 2018; Sievers et al., 2018). Their application in human cells could be particularly useful for mapping functional consequences of human genetic variation and for engineering enhanced molecular devices in a more relevant cell model. Here, we achieved very reproducible DMS data (R^2^ = 0.92) by maintaining a high >12,500x cell coverage using suspension K562 cells in a large spinner flask, and using our novel synthetic surface marker so we could separate cells with magnetic beads instead of FACS, demonstrating an approach to high-quality and lower-difficulty DMS in human cells. HT-recruit opens up the possibility of performing deep mutational scans on the hundreds of human activator and repressor domains in order to define their functional residues and better understand the biophysical requirements for effector activity.

### Applications in Synthetic Biology

Previously, a limited number of transcriptional effector domains were available for the engineering of synthetic transcription factors. HT-recruit enables the discovery of effector domains that can upregulate or downregulate transcription, and the identification of mutants of effector domains with enhanced activity. Here, we identified N-terminal substitutions that improve the expression level of the ZNF10 KRAB effector and could readily be ported into the CRISPRi system. Specifically, we identify poorly conserved, unnecessary residues that could be replaced with expression level-enhancing residues to improve the effector; this approach could be a generally applicable strategy to improve domains beyond KRAB.

The new transcriptional effector domains reported here have several advantages for applications that rely on synthetic transcription factors. We identify short domains (≤80 amino acids) and demonstrate a process for shortening them further to a minimally sufficient sequence as short at 10 amino acids, which is an advantage for delivery (e.g. packaging in viral vectors). The domains are extracted from human proteins, which is advantageous for avoiding immunogenicity associated with viral effector domains.

### High-throughput protein domain functional screens in human cells

This work is the first effort to systematically characterize transcriptional effector domains in human cells, to our knowledge. It expands the catalog of functional transcriptional effector domains but is certainly still incomplete. We envision new library designs that tile transcription factors or focus on regions with activator-like signatures will identify additional human activator domains, as such designs have uncovered activators in yeast and Drosophila experiments (Arnold et al., 2018; Erijman et al., 2020; Ravarani et al., 2018). Without major modifications, HT-recruit should be compatible with any cell type that is transfectable (to integrate the reporter) and transducible by lentivirus (to deliver the library).

Our libraries were made possible by recent increases in the length of pooled oligonucleotides. Using 300 nucleotide libraries, we were able to encode up to 80 amino acid domains and test the majority of Pfam domains in nuclear proteins. Future improvements in synthesis length can be expected to enable more complete domain libraries. Additionally, larger sequences could potentially be accessed with alternative cloning approaches, including using oligonucleotide assembly methods (Sidore et al., 2020).

We made our screens more scalable by engineering a novel synthetic cell surface marker that enables more efficient, inexpensive, and rapid screening of these libraries using magnetic separation. Magnetic separation, which we previously applied to CRISPR screens (Haney et al., 2018), makes high-throughput screening more accessible in comparison to the conventional approach of sorting libraries based on fluorescent reporter gene expression, which can be a technical bottleneck. The synthetic surface marker could facilitate the adaptation to magnetic separation for other reporter-based assays, including Massively Parallel Reporter Assays (MPRA).

More broadly, the strategy we describe here for designing and screening pooled protein domain libraries in high-throughput can readily be applied beyond transcriptional effectors.

## Supplementary Figures

**Supplementary Figure 1.**
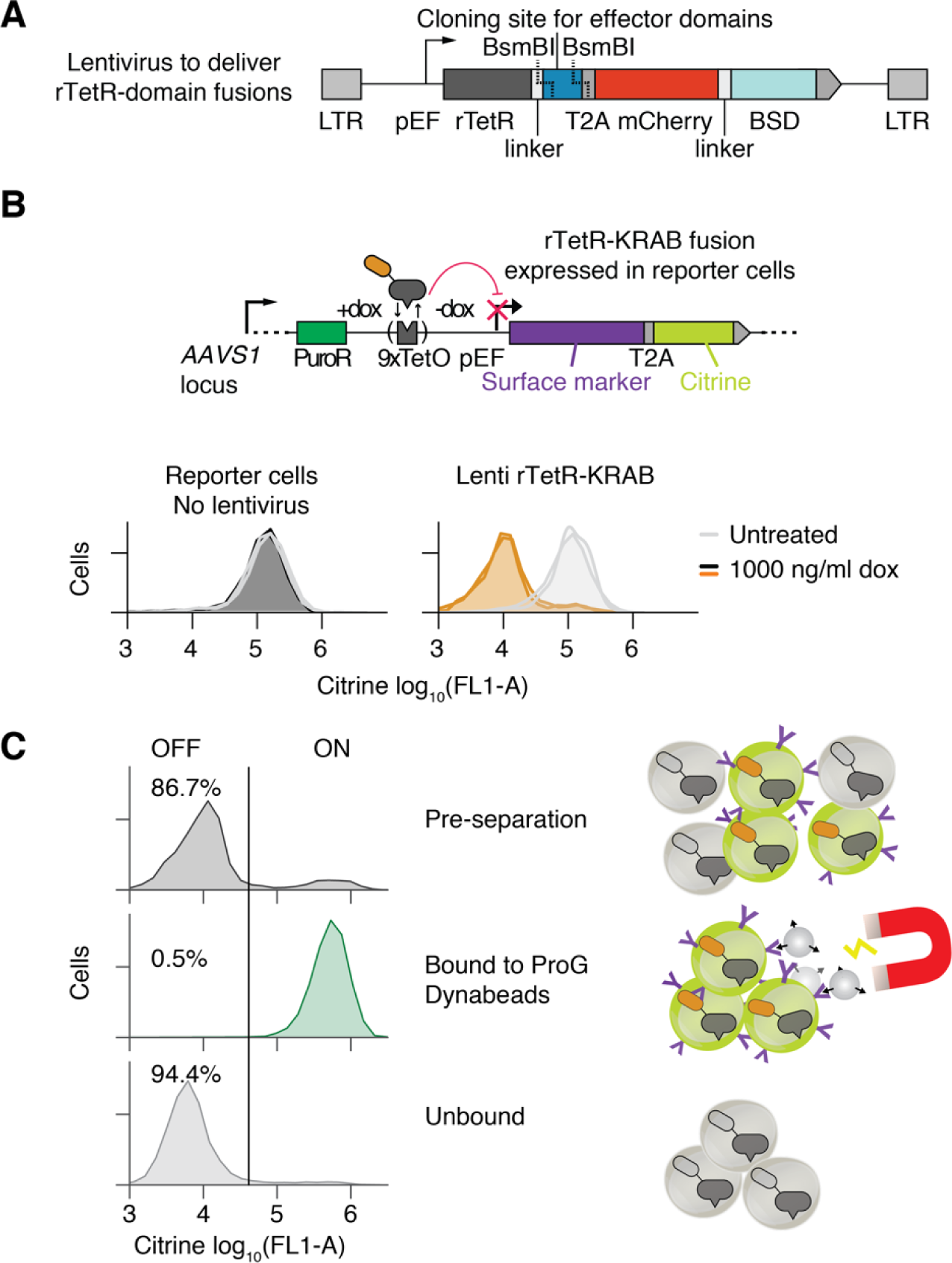
Validation of lentiviral recruitment assay and dual reporter for gene silencing. A. Schematic of lentiviral recruitment vector with Golden Gate cloning site for creating fusions of effector domains to the dox-inducible DNA-binding domain rTetR. The constitutive pEF promoter drives expression of the rTetR-effector fusion and mCherry-BSD (Blasticidin S deaminase resistance gene), separated by a T2A self-cleaving peptide. B. (Top) Schematic of rTetR-KRAB fusion recruitment to the dual reporter gene. The reporter is integrated in the *AAVS1* locus by TALEN-mediated homology-directed repair and the PuroR resistance gene is driven by the endogenous *AAVS1* promoter. The dual reporter consists of a synthetic surface marker (Igκ-hIgG1-Fc-PDGFRβ) and a citrine fluorescent protein. (Bottom) Pilot test in K562 reporter cells. Reporter cells were generated by TALEN-mediated homology-directed repair to integrate the reporter into the *AAVS1* locus and then selected with puromycin. Cells were then spinfected with lentivirus to deliver rTetR-KRAB, and then either left untreated or treated with 1000 ng/ml doxycycline to induce rTetR binding to DNA at the TetO sites. Untreated cell distributions are shown in light grey and doxycycline-treated cells are shown in black or orange, with two independently-transduced biological replicates in each condition. The lentivirus-treated cells are gated on mCherry as the delivery marker. The KRAB domain from human ZNF10 was used. C. Demonstration of magnetic separation of OFF from ON cells using ProG Dynabeads that bind to the synthetic surface marker. Ten million cells were subjected to magnetic separation using 30 µl of beads, and the citrine reporter expression was measured before and after by flow cytometry. Illustration of mixed ON and OFF cells being subjected to magnetic separation is shown on the right.

**Supplementary Figure 2.**
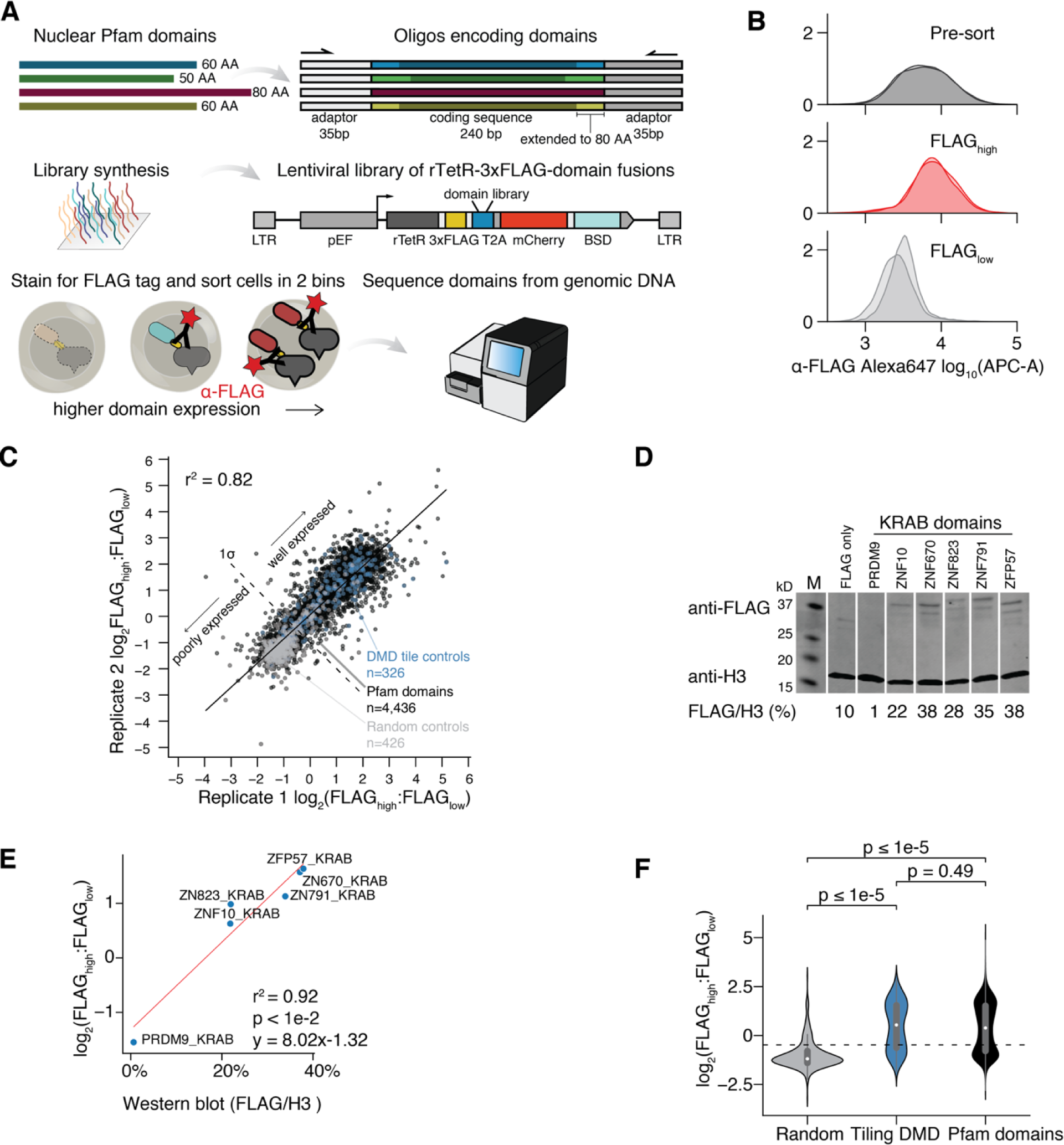
High-throughput measurements of domain expression by FLAG staining, sorting, and sequencing. A. (Top) Schematic of high-throughput strategy for measuring the expression level of each domain in the library. Domains under 80AA long are extended on both sides, using their native protein sequence, to reach 80AA so all synthesized library elements are the same length. (Middle) The library is cloned into a FLAG-tagged construct and delivered to K562 cells by lentivirus at low multiplicity of infection, such that the majority of cells express a single library member. The mCherry-BSD fusion protein enables blasticidin selection and a fluorescent marker for delivery and selection efficiency, without the use of a second 2A component. (Bottom) Expression is measured by staining the cells with anti-FLAG, sorting high and low expression populations, sequencing the domains, and computing the log_2_(FLAG_high_:FLAG_low_) ratio. B. Distribution of FLAG staining levels measured by flow cytometry before and after sorting into two bins (N = 2 biological replicates of the cell library shown with overlapping shaded areas). C. Reproducibility of biological replicates from the domain expression screen (r^2^ = 0.82). Well-expressed domains, above the threshold (dashed line one standard deviation above the median of the random controls), were selected for further analysis in the transcriptional regulation screens. D. Validation of expression level for a panel of KRAB domains. Individual rTetR-3XFLAG-KRAB constructs were delivered to K562 cells by lentivirus. Cells were selected with blasticidin and confirmed to be >80% mCherry positive by flow cytometry. Expression level was measured by Western blot with anti-FLAG antibody. Anti-histone H3 was used as a loading control for normalization. Levels were quantified using ImageJ. E. Comparison of high-throughput measurements of expression with Western blots protein levels. These 6 KRAB domains were cloned individually using the exact 80 AA sequence from the Pfam domain library. F. Distribution of expression levels for different categories of library members. Random controls are poorly expressed compared to tiles across the DMD protein or Pfam domains (p<1e-5, Mann Whitney test). Dashed line shows the threshold for expression level, as in (C).

**Supplementary Figure 3.**
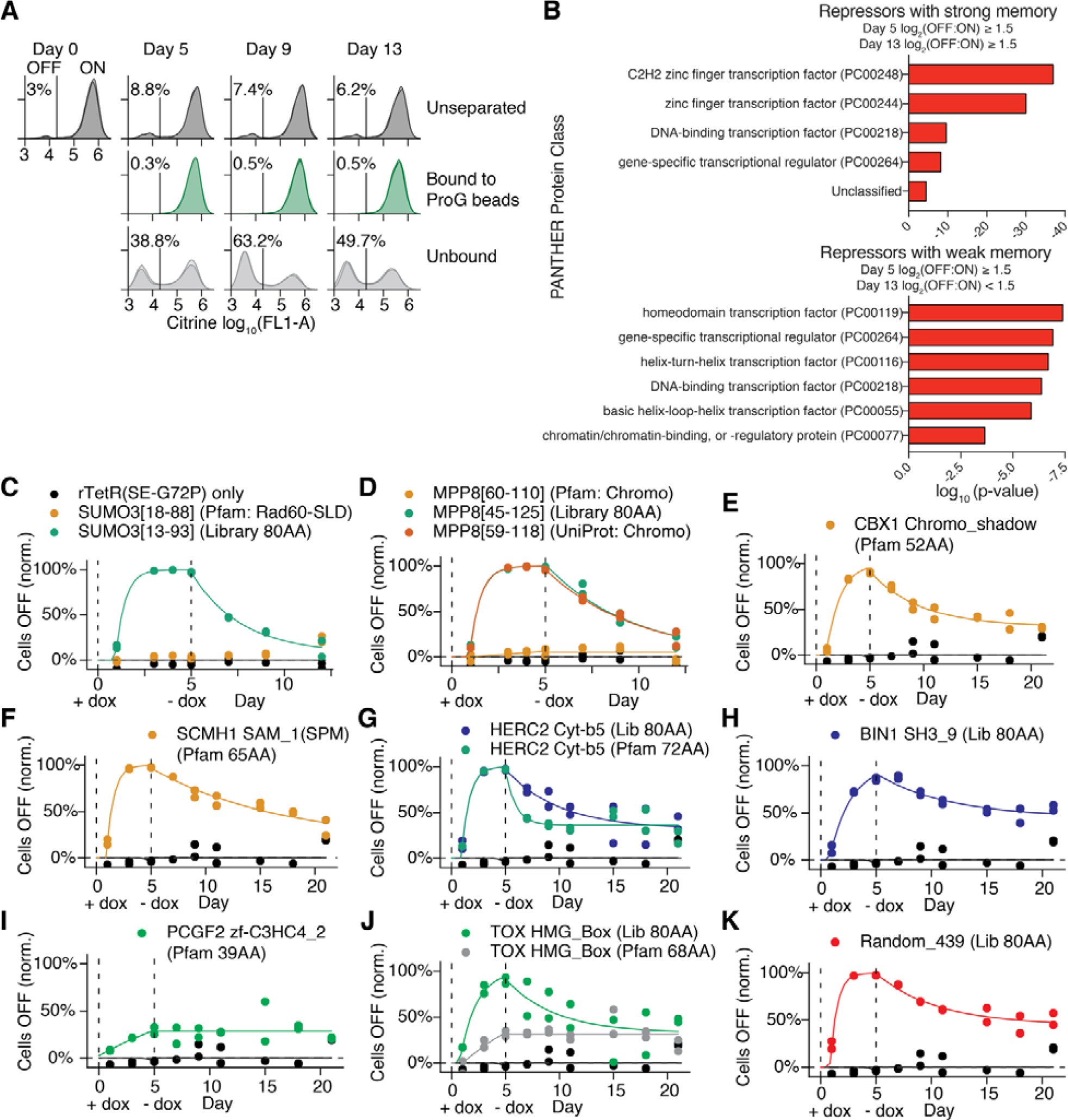
HT-recruit identifies domains with repressor function. A. Flow cytometry shows citrine reporter level distributions in the pool of cells expressing the Pfam domain library, before and after magnetic separation using ProG DynaBeads that bind to the synthetic surface marker. Overlapping histograms are shown for two biological replicates. The average percentage of cells OFF is shown to the left of the vertical line showing the citrine level gate. 1000 ng/ml doxycycline was added on Day 0 and removed on Day 5. B. PANTHER protein class enrichments for nuclear proteins that contain repressor domains with stronger or weaker memory, when compared to the background set of all nuclear proteins with domains included in our library. C. rTetR-SUMO validation time courses fit with gene silencing model (**Methods**). The 80 AA sequence centered around the Rad60-SLD domain of SUMO3 and the trimmed domain were individually cloned into lentivirus and delivered to the reporter cells. 1000 ng/ml doxycycline was added on day 0 and removed on day 5 (N = 2 biological replicates). The fraction of mCherry-positive cells with the citrine reporter OFF was determined by flow cytometry and normalized for background silencing using the untreated, time-matched controls (**Methods**). D. HUSH complex member MPP8 Chromo domain validation with the full 80 AA sequence used in the screen and sequences trimmed to match the Pfam and UniProt annotations. E. CBX1 Chromoshadow domain validation with 52 AA sequence trimmed to match the Pfam annotation. F. Polycomb 1 component SCMH1 SAM1 domain (also known as SPM) validation with 65 AA sequence trimmed to match the Pfam annotation. G. HERC2 Cyt-b5 domain validation with the full 80 AA sequence used in the screen and a 72 AA sequence trimmed to match the Pfam annotation. H. BIN1 SH3_9 domain validation. I. Polycomb 1 component PCGF2 zf-C3HC4_2 domain validation with 39 AA sequence trimmed to match the Pfam annotation. J. TOX HMG box domain validation with the full 80 AA sequence used in the screen and a 68 AA sequence trimmed to match the Pfam annotation. K. Validation of a random 80 AA sequence that functions as a repressor.

**Supplementary Figure 4.**
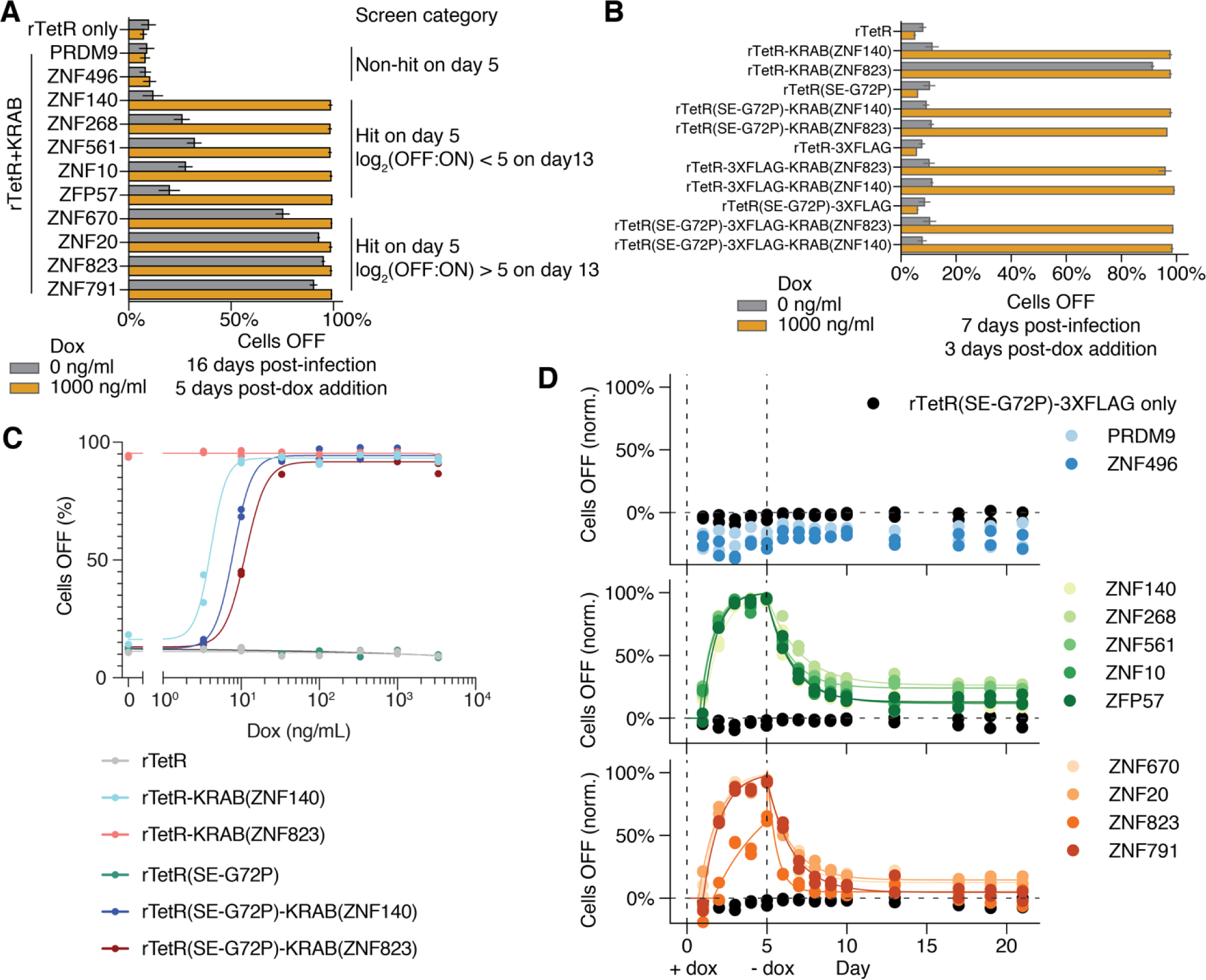
rTetR(SE-G72P) mitigates leaky KRAB silencing in human cells. A. Silencing by rTetR-KRAB fusions, showing leaky silencing without doxycycline treatment for a subset of KRAB domains (high gray bars). Constructs were delivered to reporter cells by lentivirus at day 0, cells were selected with blasticidin between days 3 and 11, cells were split into a doxycycline-treated or untreated condition at day 11, and reporter levels were measured by flow cytometry at day 16. Results are shown after gating for the mCherry positive cells. The KRAB domains were selected from three categories based on their measurements in the screen, labeled on the right. The bar shows the average and the error bar shows the standard deviation (N = 3 independently transduced biological replicates). B. Leakiness can be mitigated by using rTetR(SE-G72P) or introducing 3XFLAG between rTetR and the KRAB domain from ZNF823. Constructs were delivered to reporter cells by lentivirus at day 0, cells were split into a doxycycline-treated or untreated condition at day 4, and reporter levels were measured by flow cytometry at day 7. Results are shown after gating for the mCherry positive cells. A non-leaky KRAB domain from ZNF140 was used as a control. The bar shows the average and the error bar shows the standard deviation (N = 2 independently transduced biological replicates). C. The K562 reporter cell lines with stable lentiviral expression of either a leaky KRAB domain from ZNF823 or a non-leaky repressor KRAB domain from ZNF140 cloned as fusions with either rTetR or rTetR(SE-G72P) were treated with varied doses of doxycycline. Reporter levels were measured by flow cytometry four days later, and the percentage of mCherry positive cells with the citrine reporter OFF is shown (N = 2 independently transduced biological replicates). The dose response was fit by least squares with a non-linear variable slope sigmoidal curve using PRISM statistical analysis software. D. Silencing and memory dynamics for all individual validations of KRAB domains, fit with the gene silencing model (**Methods**). rTetR(SE-G72P)-KRAB fusions were delivered to K562 reporter cells by lentivirus, selected with blasticidin, and then 10 ng/ml doxycycline was added on day 0 and removed on day 5 (N = 2 biological replicates). The fraction of mCherry positive cells with the citrine reporter OFF was determined by flow cytometry and normalized for background silencing using the untreated, time-matched controls. We used 10 ng/ml dox to work in a dynamic range where it is easier to measure differences in silencing and memory capabilities between fast KRAB silencing domains. With 1000 ng/ml doxycycline, all of the repressor hit KRAB domains here (greens and oranges) fully silence the reporter within 5 days with indistinguishable dynamics (data not shown). Notably, the KRABs that were leaky on rTetR (oranges), do not show significantly different memory dynamics from the KRABs that were not leaky (greens) when fused to the rTetR(SE-G72P). Importantly, none of the rTetR(SE-G72P)-KRAB fusions showed significant leaky silencing in the untreated condition.

**Supplementary Figure 5.**
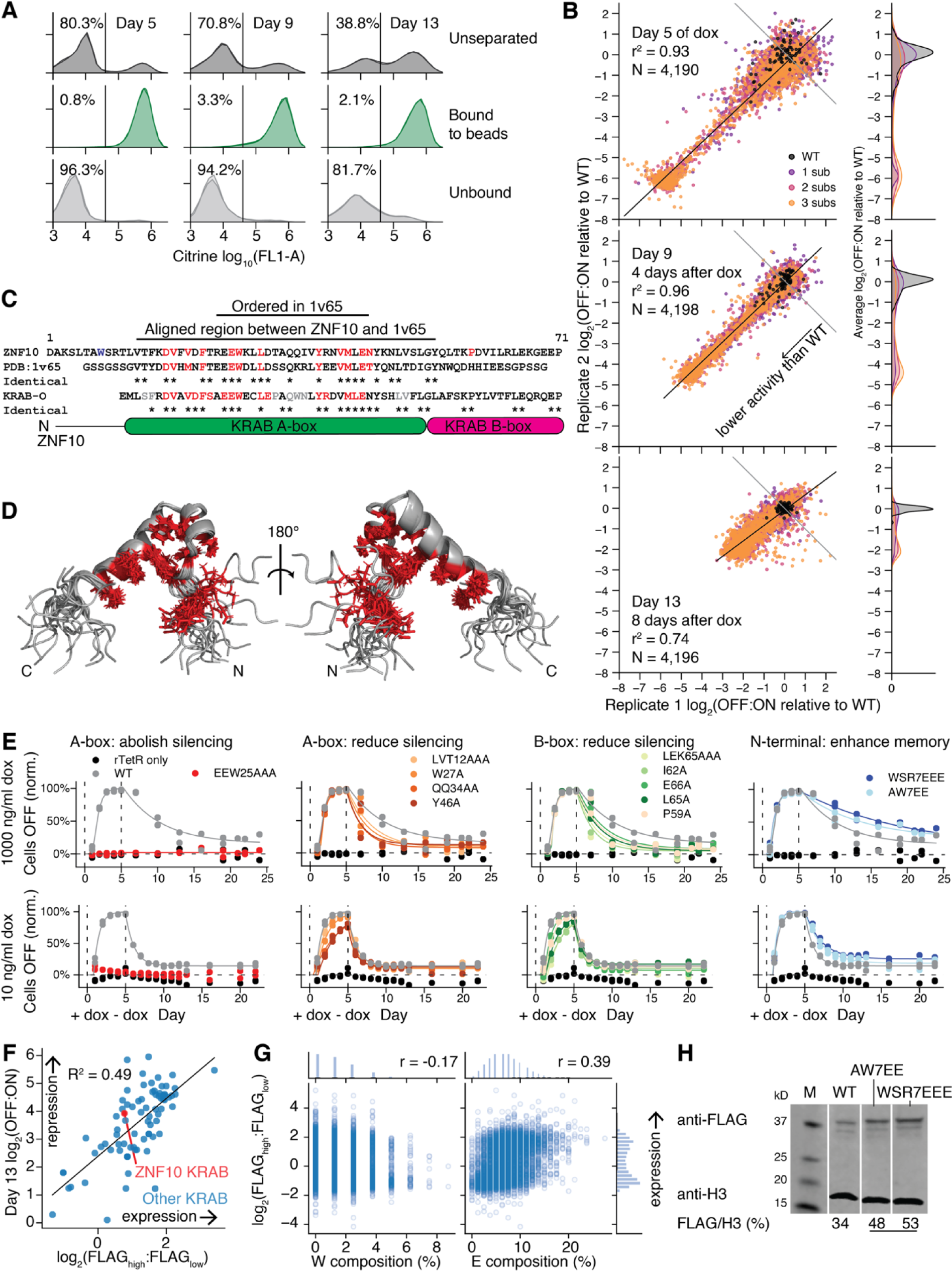
Deep mutational scan of ZNF10 KRAB used in CRISPRi. A. Flow cytometry shows citrine reporter levels in the cells with the pooled KRAB library, before and after magnetic separation using ProG DynaBeads that bind to the synthetic surface marker. Overlapping histograms are shown for two biological replicates. The average percentage of cells OFF is shown to the left of the vertical line showing the citrine level gate. B. OFF:ON enrichments from two biological replicates of the deep mutational library of the ZNF10 KRAB domain at days 5, 9 and 13. Cells were treated with 1000 ng/ml doxycycline for the first 5 days. Grey diagonal lines show where the average log_2_(OFF:ON) is the median of the WT domains (black dots). The black diagonal lines show the fit linear model. C. Alignment of human ZNF10 KRAB with mouse KRAB used in the NMR structure (PDB:1v65) and KRAB-O used in the recombinant protein binding assays (Peng et al., 2009). The ordered region is used in **Figure 3F** and the aligned region containing all 12 necessary residues is used in **Figure S5D**. The residues necessary for silencing at day 5 are colored in red in the ZNF10 and PDB:1v65 sequences. The residues necessary for binding recombinant KAP1 are colored in red and the residues unnecessary for binding KAP1 are colored in grey in the KRAB-O sequence, summarizing previously published results (Peng et al., 2009). D. Ensemble of 20 states of the KRAB NMR structure (PDB:1v65). The residues necessary for silencing at day 5 are colored in red. E. Silencing and memory dynamics for all individual validations of KRAB ZNF10 mutants, fit with the gene silencing model (**Methods**). (Top) rTetR-KRAB fusions were delivered to K562 reporter cells by lentivirus, selected with blasticidin, and then 1000 ng/ml doxycycline was added on day 0 and removed on day 5 (N = 2 biological replicates. (Bottom) rTetR(SE-G72P)-KRAB fusions were delivered to K562 reporter cells by lentivirus, selected with blasticidin, and then 10 ng/ml doxycycline was added on day 0 and removed on day 5 (N = 2 biological replicates). The column labels describe the variant location within the KRAB domain and impact on effector function. The fraction of mCherry positive cells with the citrine reporter OFF was determined by flow cytometry and normalized for background silencing using the untreated, time-matched controls. All of the rTetR(SE-G72P)-KRAB fusions were also measured over 5 days of treatment with 1000 ng/ml doxycycline and the results were indistinguishable from those with rTetR, with all KRAB variants completely silencing the reporter except the EEW25AAA variant that does not silence (data not shown). F. Correlation of rTetR-KRAB fusion expression level and day 13 silencing score, from the Pfam domain library. Only KRAB domains that were shown to interact with co-repressor KAP1 by IP/MS (Helleboid et al., 2019) are included. G. Correlations of amino acid frequency with domain expression level, across the library of Pfam domains and controls (Pearson’s r value is shown). H. Western blot for FLAG-tagged rTetR-KRAB fusions after lentiviral delivery to K562. Cells were selected for delivery with blasticidin and were confirmed to be >80% mCherry positive by flow cytometry. Expression level relative to the H3 loading control was quantified using ImageJ.

**Supplementary Figure 6.**
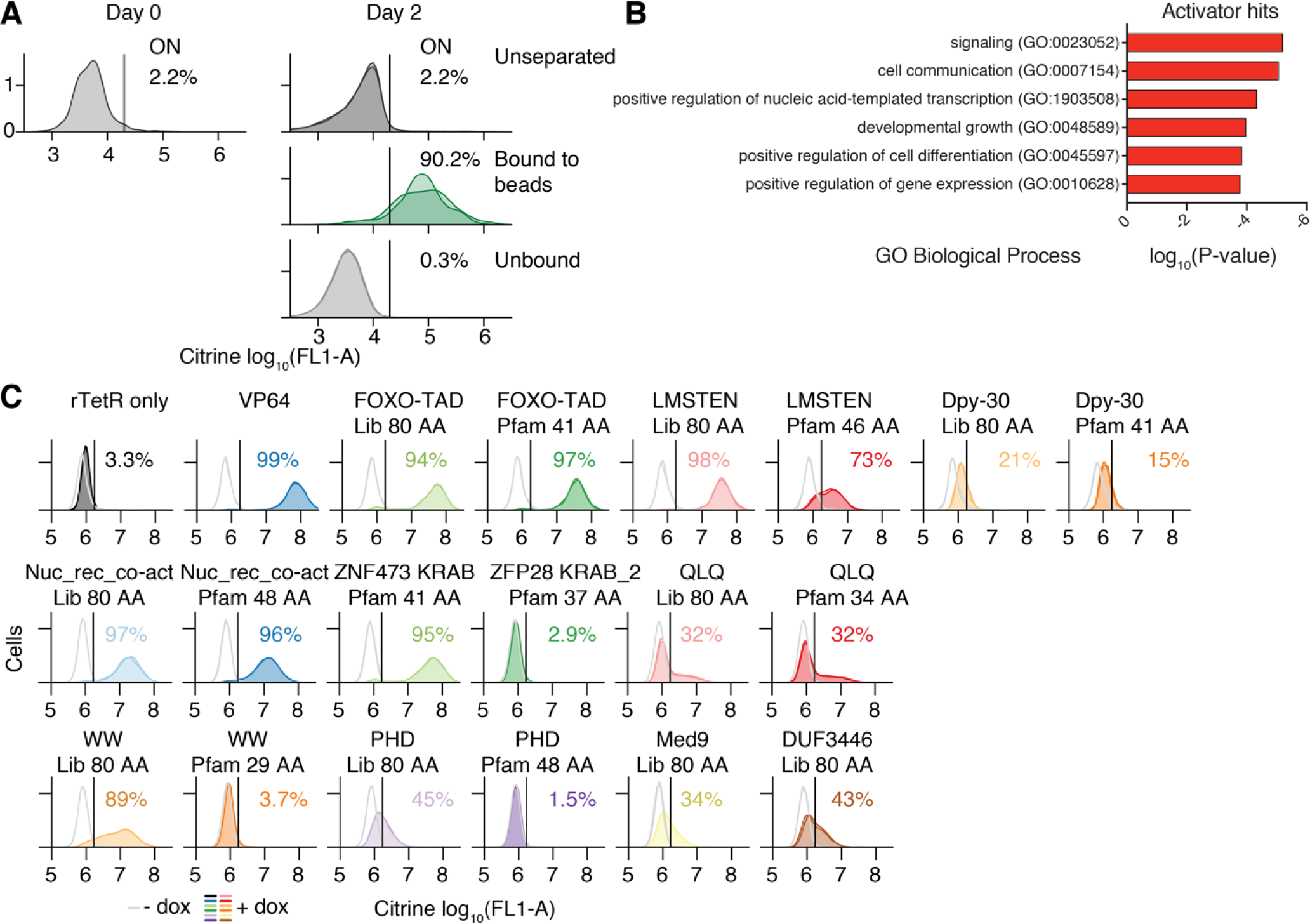
HT-recruit to a minimal promoter discovers activator domains. A. Flow cytometry for pooled library of Pfam domains in activation reporter cells, before and after magnetic separation. The percentage of cells ON is shown to the right of the citrine level gate, drawn with a vertical line. 1-2 biological replicates are shown with overlapping shaded areas. B. GO term enrichment of genes containing a hit activation domain, compared to the background set of all proteins containing a well-expressed domain in the library after counts filtering. Raw p-values are shown, and all shown terms are below a 10% false discovery rate. C. Individual validations of activator domains. rTetR(SE-G72P)-domain fusions were delivered to K562 reporter cells by lentivirus and selected with blasticidin. Cells were treated with 1000 ng/ml doxycycline for 2 days, and then citrine reporter levels were measured by flow cytometry. Untreated cell distributions are shown in light grey and doxycycline-treated cells are shown in colors, with two independently-transduced biological replicates in each condition. The vertical line shows the citrine gate used to determine the fraction of cells ON, and the average fraction ON for the doxycycline-treated cells is shown. VP64 is a positive control. Each domain was tested as both the extended 80 AA sequence from the library or the trimmed Pfam-annotated domain sequence, with the exceptions of Med9 and DUF3446 which had minimal extensions because the Pfam annotated regions are 75 and 69 AA, respectively. The corresponding results for the 80 AA library sequence for the KRAB domains are shown in **Figure 5**.

**Supplementary Figure 7.**
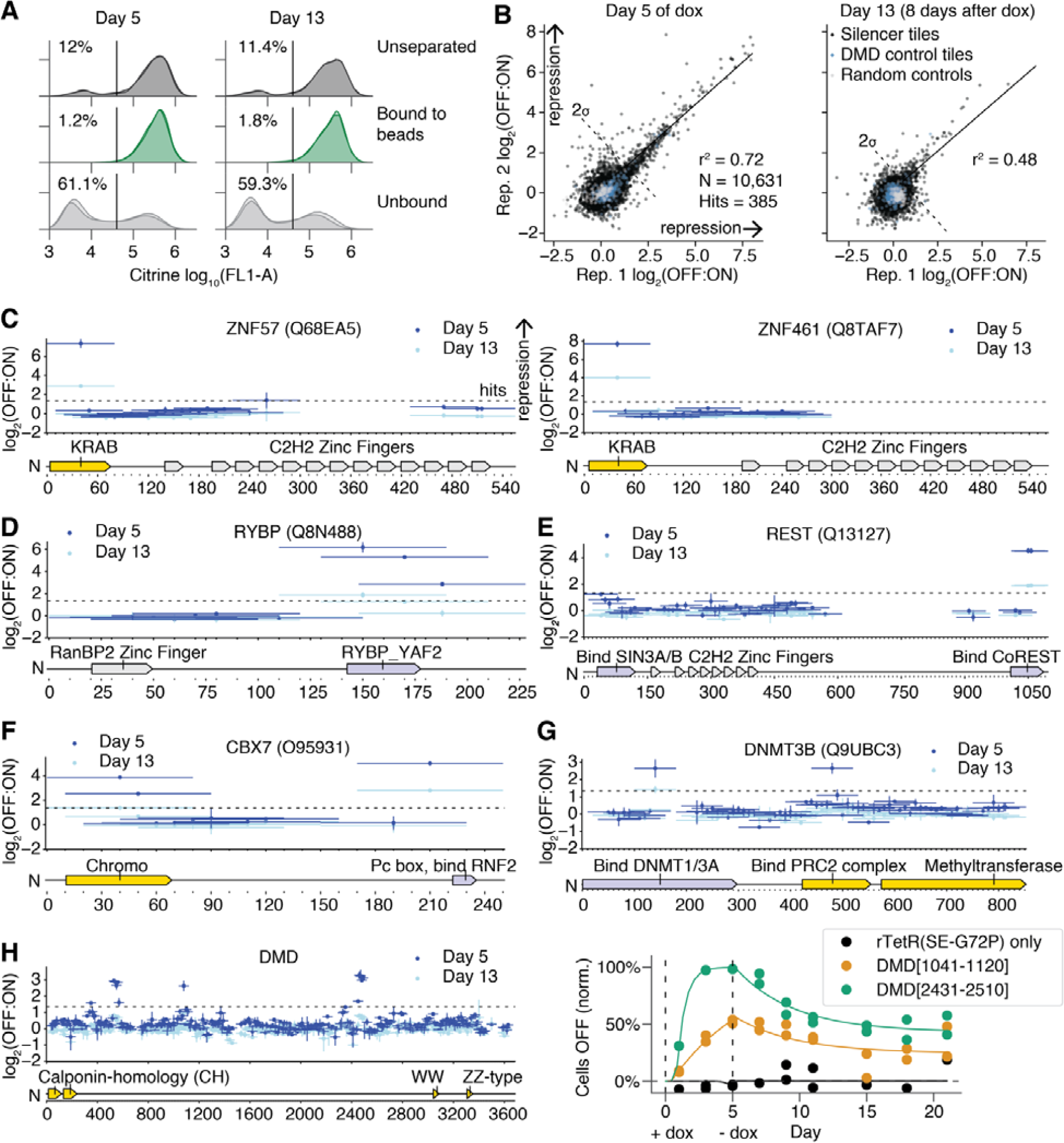
Tiling screen identifies compact repressor domains in nuclear proteins. A. Flow cytometry shows citrine reporter level distributions in the pool of cells expressing the tiling library, before and after magnetic separation using ProG DynaBeads that bind to the synthetic surface marker. Overlapping histograms are shown for two biological replicates. The average percentage of cells OFF is shown to the left of the vertical line showing the citrine level gate. 1000 ng/ml doxycycline was added on Day 0 and removed on Day 5. B. High-throughput recruitment measurements from two biological replicates of a nuclear protein tiling library at Day 5 of doxycycline treatment and Day 13, 8 days after doxycycline removal. The hit calling threshold is 2 standard deviations above the mean of the random and DMD tiling controls. C. Tiling results for KRAB Zinc finger proteins ZNF57 and ZNF461. Each bar is an 80 AA tile and the vertical error bars are the range from 2 biological replicates. Protein annotations are sourced from UniProt. D. Tiling RYBP. Diagram shows protein annotations, retrieved using the UniProt ID written at top. Vertical error bars show the standard error from two biological replicates. E. Tiling REST. F.Tiling CBX7. D. Tiling DNMT3B. H. (Left) Tiling DMD. (Right) Dynamics of silencing and memory after recruitment of DMD hit tiles. Cells were treated with 1000 ng/ml doxycycline for the first 5 days and citrine reporter levels were measured by flow cytometry. The percentage of cells OFF was normalized to account for background silencing and the data (dots) were fit with a gene silencing model (curves) (N = 2 biological replicates).

## Supplementary Tables

(available upon request from)

Supplementary Table 1: Oligonucleotide libraries used in this study.

Supplementary Table 2: Plasmids used in this study.

Supplementary Table 3: Primers used in this study.

Supplementary Table 4: HT-recruit processed data.

Supplementary Table 5: External ChIP dataset identifiers.

## Materials and Methods

### Cell lines and cell culture

All experiments presented here were carried out in K562 cells (ATCC CCL-243). Cells were cultured in a controlled humidified incubator at 37°C and 5% CO_2_, in RPMI 1640 (Gibco) media supplemented with 10% FBS (Hyclone), penicillin (10,000 I.U./mL), streptomycin (10,000 ug/mL), and L-glutamine (2 mM). HEK293FT and HEK293T-LentiX cells, used to produce lentivirus, as described below, were grown in DMEM (Gibco) media supplemented with 10% FBS (Hyclone), penicillin (10,000 I.U./mL), and streptomycin (10,000 ug/mL). Reporter cell lines were generated by TALEN-mediated homology-directed repair to integrate a donor construct into the *AAVS1* locus as follows: 1.2×10^6^ K562 cells were electroporated in Amaxa solution (Lonza Nucleofector 2b, setting T0-16) with 1000 ng of reporter donor plasmid (**Table S2**), and 500 ng of each TALEN-L (Addgene #35431) and TALEN-R (Addgene #35432) plasmid (targeting upstream and downstream the intended DNA cleavage site, respectively). After 7 days, the cells were treated with 1000 ng/mL puromycin antibiotic for 5 days to select for a population where the donor was stably integrated in the intended locus, which provides a promoter to express the PuroR resistance gene. Fluorescent reporter expression was measured by microscopy and by flow cytometry (BD Accuri).

### Nuclear protein Pfam domain library design

We queried the UniProt database (UniProt Consortium, 2015) for human genes that can localize to the nucleus. Subcellular location information on UniProt is determined from publications or ‘by similarity’ in cases where there is only a publication on a similar gene (e.g. ortholog) and is manually reviewed. We then retrieved Pfam-annotated domains using the ProDy searchPfam function (Bakan et al., 2011). We filtered for domains that were 80 amino acids or shorter and excluded the C2H2 Zinc finger DNA-binding domains, which are highly abundant, repetitive, and not expected to function as transcriptional effectors. We retrieved the sequence of the annotated domain and extended it equally on either side to reach 80 amino acids total. Duplicate sequences were removed, then codon optimization was performed for human codon usage, removing BsmBI sites and constraining GC content to between 20% and 75% in every 50 nucleotide window (performed with DNA chisel (Zulkower and Rosser, 2020)). 499 random controls of 80 amino acids lacking stop codons were computationally generated as controls. 362 elements tiling the DMD protein in 80 amino acid tiles with a 10 amino acid sliding window were also included as controls because DMD was not thought to be a transcriptional regulator. In total, the library consists of 5,955 elements (**Table S1**).

### Silencer tiling library design

216 proteins involved in transcriptional silencing were curated from a database of transcriptional regulators (Lambert et al., 2018). We manually added 32 proteins that we thought likely to be involved in transcriptional silencing and then generated an unbiased protein tiling library (**Table S1**). To do this, the canonical transcript for each gene was retrieved from the Ensembl BioMart (Kinsella et al., 2011) using the Python API. If no canonical transcript was found, the longest transcript with a CDS was retrieved. The coding sequences were divided into 80 amino acid tiles with a 10 amino acid sliding window between tiles. For each gene, a final tile was included, spanning from 80 amino acids upstream of the last residue to that last residue, such that the C-terminal region would be included in the library. Duplicate protein sequences were removed, and codon optimization was performed for human codon usage, removing BsmBI sites and constraining GC content to between 20% and 75% in every 50 nucleotide window (performed with DNA chisel (Zulkower and Rosser, 2020)). 361 DMD tiling negative controls were included, as in the previous library design, resulting in 15,737 library elements in total.

### KRAB deep mutational scan library design

A deep mutational scan of ZNF10 KRAB domain sequence, as used in CRISPRi (Gilbert et al., 2014), was designed with all possible single substitutions and all consecutive double and triple substitutions of the same amino acid (e.g. substitution with AAA) (**Table S1**). These amino acid sequences were reverse translated into DNA sequences using a probabilistic codon optimization algorithm, such that each DNA sequence contains some variation beyond the substituted residues, which improves the ability to unambiguously align sequencing reads to unique library members. In addition, all Pfam-annotated KRAB domains from human KRAB genes found on InterPro were included, similarly as in the previous nuclear Pfam domain library. Tiling sequences, as designed in the previous tiling library, were also included for five KRAB Zinc Finger genes. 300 random control sequences and 200 tiles from the DMD gene were included as negative controls. During codon optimization, BsmBI sites were removed and GC content was constrained to be between 30% and 70% in every 80 nucleotide window (performed with DNA chisel (Zulkower and Rosser, 2020)). The total library size was 5,731 elements.

### Library cloning

Oligonucleotides with lengths up to 300 nucleotides were synthesized as pooled libraries (Twist Biosciences) and then PCR amplified. 6x 50 ul reactions were set up in a clean PCR hood to avoid amplifying contaminating DNA. For each reaction, we used 5 ng of template, 0.1 µl of each 100 µM primer, 1 µl of Herculase II polymerase (Agilent), 1 µl of DMSO, 1 µl of 10 nM dNTPs, and 10 µl of 5x Herculase buffer. The thermocycling protocol was 3 minutes at 98°C, then cycles of 98°C for 20 seconds, 61°C for 20 seconds, 72°C for 30 seconds, and then a final step of 72°C for 3 minutes. The default cycle number was 29x, and this was optimized for each library to find the lowest cycle that resulted in a clean visible product for gel extraction (in practice, 25 cycles was the minimum). After PCR, the resulting dsDNA libraries were gel extracted by loading ≥4 lanes of a 2% TBE gel, excising the band at the expected length (around 300 bp), and using a QIAgen gel extraction kit. The libraries were cloned into a lentiviral recruitment vector pJT050 (Table S2) with 4×10 µl GoldenGate reactions (75 ng of pre-digested and gel-extracted backbone plasmid, 5 ng of library, 0.13 µl of T4 DNA ligase (NEB, 20000 U/µl), 0.75 µl of Esp3I-HF (NEB), and 1 µl of 10x T4 DNA ligase buffer) with 30 cycles of digestion at 37°C and ligation at 16°C for 5 minutes each, followed by a final 5 minute digestion at 37°C and then 20 minutes of heat inactivation at 70°C. The reactions were then pooled and purified with MinElute columns (QIAgen), eluting in 6 ul of ddH_2_O. 2 µl per tube was transformed into two tubes of 50 µl of electrocompetent cells (Lucigen DUO) following the manufacturer’s instructions. After recovery, the cells were plated on 3 - 7 large 10” x 10” LB plates with carbenicillin. After overnight growth at 37°C, the bacterial colonies were scraped into a collection bottle and plasmid pools were extracted with a HiSpeed Plasmid Maxiprep kit (QIAgen). 2 - 3 small plates were prepared in parallel with diluted transformed cells in order to count colonies and confirm the transformation efficiency was sufficient to maintain at least 30x library coverage. To determine the quality of the libraries, the domains were amplified from the plasmid pool and from the original oligo pool by PCR with primers with extensions that include Illumina adapters (**Table S3**) and sequenced. The PCR and sequencing protocol were the same as described below for sequencing from genomic DNA, except these PCRs use 10 ng of input DNA and 17 cycles. These sequencing datasets were analyzed as described below to determine the uniformity of coverage and synthesis quality of the libraries. In addition, 20 - 30 colonies from the transformations were Sanger sequenced (Quintara) to estimate the cloning efficiency and the proportion of empty backbone plasmids in the pools.

### High-throughput recruitment to measure repressor activity

Large scale lentivirus production and spinfection of K562 cells were performed as follows: To generate sufficient lentivirus to infect the libraries into K562 cells, we plated HEK293T cells on four 15-cm tissue culture plates. On each plate, 9×10^5^ HEK293T cells were plated in 30 mL of DMEM, grown overnight, and then transfected with 8 μg of an equimolar mixture of the three third-generation packaging plasmids and 8 μg of rTetR-domain library vectors using 50 μl of polyethylenimine (PEI, Polysciences #23966). After 48 hours and 72 hours of incubation, lentivirus was harvested. We filtered the pooled lentivirus through a 0.45-μm PVDF filter (Millipore) to remove any cellular debris. For the nuclear Pfam domain repressor screen, 4.5×10^7^ K562 reporter cells were infected with the lentiviral library by spinfection for 2 hours, with two separate biological replicates of the infection. Infected cells grew for 3 days and then the cells were selected with blasticidin (10 μg/mL, Sigma). Infection and selection efficiency were monitored each day using flow cytometry to measure mCherry (BD Accuri C6). Cells were maintained in spinner flasks in log growth conditions each day by diluting cell concentrations back to a 5×10^5^ cells/mL, with at least 1.5×10^8^ cells total remaining per replicate such that the lowest maintenance coverage was >25,000× cells per library element (a very high coverage level that compensates for losses from incomplete blasticidin selection, library preparation, and library synthesis errors). On day 6 post-infection, recruitment was induced by treating the cells with 1000 ng/ml doxycycline (Fisher Scientific) for 5 days, then cells were spun down out of doxycycline and blasticidin and maintained in untreated RPMI media for 8 more days, up to Day 13 counting from the addition of doxycycline. 2.5×10^8^ cells were taken for measurements at each timepoint (days 5, 9, and 13). The protocol was similar for the KRAB DMS, but doxycycline was added on day 8 post-infection, >12,500× coverage, and 2×10^8^ - 2.2×10^8^ cells were taken for each timepoint. The protocol was similar for the tiling screen, but 9.6×10^7^ cells were infected, doxycycline was added on day 8 post-infection, at least 2×10^8^ cells were maintained at each passage for >12,500× coverage, and 2×10^8^ - 2.7×10^8^ cells were taken for each timepoint.

### High-throughput recruitment to measure transcriptional activation activity

For the nuclear Pfam domain activator screen, lentivirus for the nuclear Pfam library in the rTetR(SE-G72P)-3XFLAG vector (**Table S2**) was generated as for the repressor screen, and 3.8×10^7^ K562-pDY32 minCMV reporter cells (**Table S2**) were infected with the lentiviral library by spinfection for 2 hours, with two separate biological replicates of the infection. Infected cells grew for 2 days and then the cells were selected with blasticidin (10 μg/mL, Sigma). Infection and selection efficiency were monitored each day using flow cytometry to measure mCherry (BD Accuri C6). Cells were maintained in spinner flasks in log growth conditions each day by diluting cell concentrations back to a 5×10^5^ cells/mL, with at least 1×10^8^ total cells remaining per replicate such that the lowest maintenance coverage was >18,000× cells per library element. On day 7 post-infection, recruitment was induced by treating the cells with 1000 ng/ml doxycycline (Fisher Scientific) for 2 days, then cells were spun down out of doxycycline and blasticidin and maintained in untreated RPMI media for 4 more days. 2×10^8^ cells were taken for measurements at the day 2 time point. There was no evidence of activation memory at day 4 post-doxycycline removal, as determined by the absence of citrine positive cells by flow cytometry, so no additional time points were collected.

### Magnetic separation of reporter cells

At each timepoint, cells were spun down at 300 × *g* for 5 minutes and media was aspirated. Cells were then resuspended in the same volume of PBS (Gibco) and the spin down and aspiration was repeated, to wash the cells and remove any IgG from serum. Dynabeads(tm) M-280 Protein G (ThermoFisher 10003D) were resuspended by vortexing for 30 seconds. 50 mL of blocking buffer was prepared per 2×10^8^ cells by adding 1 gram of biotin-free BSA (Sigma Aldrich) and 200 μl of 0.5 M pH 8.0 EDTA (ThemoFisher 15575020) into DPBS (Gibco), vacuum filtering with a 0.22-μm filter (Millipore), and then kept on ice. 60 μl of beads was prepared for every 1×10^7^ cells, by adding 1 mL of buffer per 200 μl of beads, vortexing for 5 seconds, placing on a magnetic tube rack (Eppendorf), waiting one minute, removing supernatant, and finally removing the beads from the magnet and resuspending in 100 - 600 μl of blocking buffer per initial 60 μl of beads. For the KRAB DMS only, 30 μl of beads was prepared for every 1×10^7^ cells, in the same way. Beads were added to cells at no more than 1×10^7^ cells per 100 μl of resuspended beads, and then incubated at room temperature while rocking for 30 minutes. For a sample with 2×10^8^ cells, we used 1.2 mL of beads, resuspended in 12 mL of blocking buffer, in a 15 mL Falcon tube and a large magnetic rack. For a sample with <5×10^7^ cells, we used non-stick Ambion 1.5 mL tubes and a small magnetic rack. After incubation, the bead and cell mixture were placed on the magnetic rack for >2 minutes. The unbound supernatant was transferred to a new tube, placed on the magnet again for >2 minutes to remove any remaining beads, and then the supernatant was transferred and saved as the unbound fraction. Then, the beads were resuspended in the same volume of blocking buffer, magnetically separated again, the supernatant was discarded, and the tube with the beads was kept as the bound fraction. The bound fraction was resuspended in blocking buffer or PBS to dilute the cells (the unbound fraction is already dilute). Flow cytometry (BD Accuri) was performed using a small portion of each fraction to estimate the number of cells in each fraction (to ensure library coverage was maintained) and to confirm separation based on citrine reporter levels (the bound fraction should be >90% citrine positive, while the unbound fraction is more variable depending on the initial distribution of reporter levels). Finally, the samples were spun down and the pellets were frozen at -20°C until genomic DNA extraction.

### High-throughput measurement of domain fusion protein expression level

The expression level measurements were made in K562-pDY32 cells (with citrine OFF) infected with the 3XFLAG-tagged nuclear Pfam domain library. 1×10^8^ cells per biological replicate were used after 5 days of blasticidin selection (10 μg/mL, Sigma), which was 7 days post-infection. 1×10^6^ control K562-JT039 cells (citrine ON, no lentiviral infection) were spiked into each replicate. Fix Buffer I (BD Biosciences, BDB557870) was preheated to 37°C for 15 minutes and Permeabilization Buffer III (BD Biosciences, BDB558050) and PBS (Gibco) with 10% FBS (Hyclone) were chilled on ice. The library of cells expressing domains was collected and cell density was counted by flow cytometry (BD Accuri). To fix, cells were resuspended in a volume of Fix Buffer I (BD Biosciences, BDB557870) corresponding to pellet volume, with 20 μl per 1 million cells, at 37°C for 10 - 15 minutes. Cells were washed with 1 mL of cold PBS containing 10% FBS, spun down at 500 × *g* for 5 minutes and then supernatant was aspirated. Cells were permeabilized for 30 minutes on ice using cold BD Permeabilization Buffer III (BD Biosciences, BDB558050), with 20 μl per 1 million cells, which was added slowly and mixed by vortexing. Cells were then washed twice in 1 ml PBS+10% FBS, as before, and then supernatant was aspirated. Antibody staining was performed for 1 hour at room temperature, protected from light, using 5 μl / 1×10^6^ cells of α-FLAG-Alexa647 (RNDsystems, IC8529R). We then washed the cells and resuspended them at a concentration of 3×10^7^ cells / ml in PBS+10%FBS. Cells were sorted into two bins based on the level of APC-A fluorescence (Sony SH800S) after gating for mCherry positive viable cells. A small number of unstained control cells was also analyzed on the sorter to confirm staining was above background. The spike-in citrine positive cells were used to assess the background level of staining in cells known to lack the 3XFLAG tag, and the gate for sorting was drawn above that level. After sorting, the cellular coverage ranged from 336 - 1,295 cells per library element across samples. The sorted cells were spun down at 500 × *g* for 5 minutes and then resuspended in PBS. Genomic DNA extraction was performed following the manufacturer’s instructions (QIAgen Blood Maxi kit was used for samples with >1×10^7^ cells, and QIAamp DNA Mini kit with one column per up to 5×10^6^ cells was used for samples with ≤1×10^7^ cells) with one modification: the Proteinase K + AL buffer incubation was performed overnight at 56°C.

### Library preparation and sequencing

Genomic DNA was extracted with the QIAgen Blood Maxi Kit following the manufacturer’s instructions with up to 1.25×10^8^ cells per column. DNA was eluted in EB and not AE to avoid subsequence PCR inhibition. The domain sequences were amplified by PCR with primers containing Illumina adapters as extensions (**Table S3**).

A test PCR was performed using 5 μg of genomic DNA in a 50 μl (half-size) reaction to verify if the PCR conditions would result in a visible band at the expected size for each sample. Then, 12 - 24x 100 μl reactions were set up on ice (in a clean PCR hood to avoid amplifying contaminating DNA), with the number of reactions depending on the amount of genomic DNA available in each experiment. 10 μg of genomic DNA, 0.5 μl of each 100 μM primer, and 50 μl of NEBnext 2x Master Mix (NEB) was used in each reaction. The thermocycling protocol was to preheat the thermocycler to 98°C, then add samples for 3 minutes at 98°C, then 32x cycles of 98°C for 10 seconds, 63°C for 30 seconds, 72°C for 30 seconds, and then a final step of 72°C for 2 minutes. All subsequent steps were performed outside the PCR hood. The PCR reactions were pooled and ≥140 μl were run on at least three lanes of a 2% TBE gel alongside a 100-bp ladder for at least one hour, the library band around 395 bp was cut out, and DNA was purified using the QIAquick Gel Extraction kit (QIAgen) with a 30 ul elution into non-stick tubes (Ambion). A confirmatory gel was run to verify that small products were removed. These libraries were then quantified with a Qubit HS kit (Thermo Fisher), pooled with 15% PhiX control (Illumina), and sequenced on an Illumina NextSeq with a High output kit using a single end forward read (266 or 300 cycles) and 8 cycle index reads.

### Domain sequencing analysis

Sequencing reads were demultiplexed using bcl2fastq (Illumina). A Bowtie reference was generated using the designed library sequences with the script ‘makeIndices.py’ and reads were aligned with 0 mismatch allowance using the script ‘makeCounts.py’. The enrichments for each domain between OFF and ON (or FLAG_high_ and FLAG_low_) samples were computed using the script ‘makeRhos.py’. Domains with <5 reads in both samples for a given replicate were dropped from that replicate (assigned 0 counts), whereas domains with <5 reads in one sample would have those reads adjusted to 5 in order to avoid the inflation of enrichment values from low depth. For all of the nuclear domain screens, domains with ≤ 5 counts in both replicates of a given condition were filtered out of downstream analysis. For the nuclear domain expression screen, well-expressed domains were those with a log_2_(FLAG_high_:FLAG_low_) ≥ 1 standard deviation above the median of the random controls. For the nuclear Pfam domain repressor screen, hits were domains with log_2_(OFF:ON) ≥ 2 standard deviations above the mean of the poorly expressed domains. For the nuclear domain activator screen, hits were domains with log_2_(OFF:ON) ≤ 2 standard deviations below the mean of the poorly expressed domains. For the silencer tiling screen, tiles with ≤20 counts in both replicates of a given condition were filtered out and hits were tiles with log_2_(OFF:ON) ≥ 2 standard deviations above the mean of the random and DMD tiling controls. Gene ontology analysis enrichments were computed using the PantherDB web tool (www.pantherdb.org). The background sets were all proteins containing domains that were well-expressed and measured in the experiment after count filters were applied. P-values for statistical significance were calculated using Fisher’s exact test, the False Discovery Rate (FDR) was computed, and only the most significant results, all with FDR<10%, were shown.

### Western blot and co-immunoprecipitation

K562 reporter cells were transduced with a lentiviral vector containing an rTetR-3XFLAG-effector-T2A-mCherry-BSD and then selected with blasticidin (10 μg/mL) until >80% of the cells were mCherry positive. 5-10 million cells were lysed in lysis buffer (1% Triton X-100, 150mM NaCl, 50mM Tris pH 7.5, Protease inhibitor cocktail). Protein amounts were quantified using the Pierce BCA Protein Assay kit (Bio-Rad). Equal amounts were loaded onto a gel and transferred to a PVDF membrane. Membrane was probed using FLAG M2 monoclonal antibody (1:1000, mouse, Sigma-Aldrich, catalog number F1804) and Histone 3 antibody (1:1000, mouse, Abcam cat no. AB1791) as primary antibodies. Goat anti-mouse IRDye 680 RD and goat anti-rabbit IRDye 800CW (1:20,000 dilution, LI-COR Biosciences, cat nos. 926-68070 and 926-32211, respectively) were used as secondary antibodies. Blots were imaged on a LiCor Odyssey CLx. Band intensities were quantified using ImageJ (Rueden et al., 2017).

### Individual repressor recruitment assays

Individual effector domains were cloned as fusions with rTetR or rTetR(SE-G72P) with or without a 3XFLAG tag (see figure legends), upstream of a T2A-mCherry-BSD marker using GoldenGate cloning into backbones pJT050 or pJT126 (**Table S2**). K562-pJT039-pEF-citrine reporter cells (**Table S2**) were then transduced with this lentiviral vector and, 3 days later, selected with blasticidin (10 μg/mL) until >80% of the cells were mCherry positive (6-7 days). Cells were split into separate wells of a 24-well plate and either treated with doxycycline (Fisher Scientific) or left untreated. After 5 days of treatment, doxycycline was removed by spinning down the cells, replacing media with DPBS (Gibco) to dilute any remaining doxycycline, and then spinning down the cells again and transferring them to fresh media. Timepoints were measured every 2-3 days by flow cytometry analysis of >7,000 cells (either a BD Accuri C6 or Beckman Coulter CytoFLEX). Data was analyzed using Cytoflow (https://github.com/bpteague/cytoflow) and custom Python scripts. Events were gated for viability and for mCherry as a delivery marker. To compute a fraction of OFF cells during doxycycline treatment, we fit a 2 component Gaussian mixture model to the untreated rTetR-only negative control cells which fits both the ON peak and the subpopulation of background-silenced OFF cells, and then set a threshold that was 2 standard deviations below the mean of the ON peak in order to label cells that have silenced as OFF. Using the time-matched untreated control, we calculated the background normalized percentage of cells *Cell*_*OFF,normalized*_ = *Cell*_*OFF*,+*dox*_ / (1 – *Cell*_*OFF,untreated*_). Two independently transduced biological replicates were used. A gene silencing model, consisting of the increasing form of the exponential decay (i.e. exponential decay subtracted from 1) during the doxycycline treatment phase and an exponential decay during the doxycycline removal phase with additional parameters for lag times before silencing and reactivation initiate, was fit to the normalized data using SciPy.

### Individual activator recruitment assays

Domains were cloned as a fusion with rTetR(SE-G72P) upstream of a T2A-mCherry-BSD marker, using GoldenGate cloning in the backbone pJT126 (**Table S2**). K562 pDY32 minCMV citrine reporter cells (**Table S2**) were then transduced with each lentiviral vector and, 3 days later, selected with blasticidin (10 μg/mL) until >80% of the cells were mCherry positive (6-7 days). Cells were split into separate wells of a 24-well plate and either treated with doxycycline or left untreated. Timepoints were measured by flow cytometry analysis of >15,000 cells (Biorad ZE5). To compute a fraction of ON cells during doxycycline treatment, we fit a Gaussian model to the untreated rTetR-only negative control cells which fits the OFF peak, and then set a threshold that was 2 standard deviations above the mean of the OFF peak in order to label cells that have activated as ON. Two independently transduced biological replicates were used.

### Analysis of amino acid residue conservation

The ZNF10 KRAB and MGA protein sequences were submitted to the ConSurf web server (Ashkenazy et al., 2010) and analyzed using the ConSeq method (Berezin et al., 2004). Briefly, ConSeq selects up to 150 homologs for a multiple string alignment, by sampling from the list of homologs with 35-95% sequence identity. Then, a phylogenetic tree is re-constructed and conservation is scored using Rate4Site (Pupko et al., 2002). ConSurf provides normalized scores, so that the average score for all residues is zero, and the standard deviation is one. The conservation scores calculated by ConSurf are a relative measure of evolutionary conservation at each residue in the protein and the lowest score represents the most conserved position in the protein. The uniqueness of the ZNF10 KRAB N-terminal extension was determined by protein BLAST to all human proteins and searching for other zinc finger protein among the BLAST matches (Johnson et al., 2008).

### Phylogenetic and alignment analyses

KRAB and homeodomain sequences were retrieved from Pfam and extended, using surrounding native sequence, to reach 80 AA. Well-expressed domains were selected for alignment. Phylogenetic trees and sequence alignments were obtained using the alignment website Clustal Omega using default parameters (McWilliam et al., 2013; Sievers et al., 2011), and the phylogenetic neighbor-joining tree without distance corrections was built with default parameters in Jalview (Waterhouse et al., 2009). Alignment visualization was performed in Jalview.

### ChIP-seq and ChIP-exo analysis

External ChIP datasets (**Table S5**) were retrieved from multiple sources. ENCODE ChIP-seq data was processed with the uniform processing pipeline of ENCODE (ENCODE Project Consortium et al., 2020), (https://www.encodeproject.org/data-standards/chip-seq/) and narrow peaks below IDR threshold 0.05 were retrieved. KRAB ZNF ChIP-exo data from tagged KRAB ZNF overexpression in HEK293 cells and KAP1 ChIP-exo data from H1 hESCs was obtained from GEO accession GSE78099 (Imbeault et al., 2017). Reads were trimmed to a uniform length of 36 basepairs and mapped to the hg38 version of the human genome using Bowtie (version 1.0.1; (Langmead et al., 2009)), allowing for up to 2 mismatches and only retaining unique alignments. Peak were called using MACS2 (version 2.1.0) (Feng et al., 2012) with the following settings: “-g hs -f BAM --keep-dup all --shift -75 --extsize 150 --nomodel”. Browser tracks were generated using custom-written Python scripts (https://github.com/georgimarinov/GeorgiScripts). For some KRAB ZNFs where ChIP-exo data was not available, ChIP-seq data from tagged KRAB ZNF overexpression in HEK293 cells was obtained from GEO accessions GSE76496 (Schmitges et al., 2016) and GSE52523 (Najafabadi et al., 2015). KRAB ZNF peaks were defined as solo binding sites if no other KRAB ZNF in the dataset had a peak less than 250 basepairs away. ENCODE H3K27ac ChIP-seq datasets for H1 cells were processed with the ENCODE pipeline (ENCODE Project Consortium et al., 2020), narrow peaks were called with MACS2, and peaks below IDR threshold 0.05 were retrieved.

### External datasets

ChIP-seq and ChIP-exo data for KRAB ZNF, KAP1, and H3K27ac (ENCODE Project Consortium et al., 2020; Imbeault et al., 2017; Najafabadi et al., 2015; Schmitges et al., 2016), KRAB ZNF gene evolutionary age (Imbeault et al., 2017), KRAB ZNF protein co-immunoprecipitation/mass-spectrometry data (Helleboid et al., 2019), and CAT assays for KRAB repressor activity (Margolin et al., 1994; Witzgall et al., 1994) were retrieved from previously published studies. The ChIP dataset identifiers are listed in **Table S5**.

## Code availability

The scripts for processing HT-recruit data and high-throughput stability assay data will be made available in the GitHub repository found at https://github.com/JoshTycko/HTRecruit.

## Data availability

We will submit the Illumina sequencing datasets to accessible online repositories.

## Material availability

We will submit the plasmids to Addgene. Requests for resources, reagents, and further information should be directed to the corresponding authors.

## Acknowledgements

We thank David Morgens, Connor Ludwig, Sarah Lensch, Roarke Kamber, William J. Greenleaf, Alistair Boettiger and members of our laboratories for helpful conversations and assistance, and Stanley Qi for allowing us to use his lab’s Cytoflex flow cytometer. We thank TWIST Biosciences for oligonucleotide library synthesis and helpful discussion. J.T. was supported by the NSF GRFP (DGE-1656518) and is supported by the NIDDK F99/K00 fellowship of the National Institutes of Health (F99DK126120). M.C.B. is supported by a grant from Stanford ChEM-H and an NIH Director’s New Innovator Award (1DP2HD08406901). L.B. is supported by a BWF-CASI award. This work was supported by a grant from NIH/ENCODE 5UM1HG009436-02 (A.K. and M.C.B.) and NIH/NIGMS R35M128947 (L.B.).

## Author contributions

J.T., M.C.B. and L.B. designed the study. J.T. designed domain libraries with contributions from M.C.B and L.B. G.T.H. and J.T. developed the magnetic separation strategy with contributions from B.K.E., A., and M.C.B. J.T., N.D., A., G.T.H., A.M., B.K.E., M.V.V., and P.S. performed experiments. J.T. analyzed data with assistance from N.D., A.M., A.K., M.C.B., and L.B. J.T., K.S., N.D., A., B.K.E., and D.Y. generated libraries, plasmids, and cell lines. G.K.M. processed ChIP-exo data and A.B. analyzed ChIP data with J.T. and A.K. J.T., N.D., M.C.B., and L.B. wrote the manuscript with contributions from all authors. M.C.B. and L.B. supervised the project.

